# Leukemic Stem Cell Subtypes Drive Distinct Niche Remodeling in Acute Myeloid Leukemia

**DOI:** 10.64898/2026.08.24.746513

**Authors:** Karin D. Prummel, Anna Mathioudaki, Ivan Berest, Shubhankar Sood, Lixiazi He, Thorsten Richter, Yaǧmur Başkan, Jonas Rauchhaus, Clarissa Holitsch, Aryan Kamal, Juan Jauregui-Lozano, Dirk Hart, Rim Moussa, Robert Reinhardt, Swati Garg, Claudia Waskow, Carsten Müller-Tidow, Sinem K. Saka, Konstantinos Kokkaliaris, Marieke A.G. Essers, Caroline Pabst, Judith B. Zaugg

**Author notes:** these authors contributed equally (Karin D. Prummel, Anna Mathioudaki, Ivan Berest). co-correspondence to and.

## Abstract

Leukemic stem cells (LSCs) sustain acute myeloid leukemia (AML) and are implicated in therapy resistance and relapse. Yet, it remains unknown how LSCs remodel the bone marrow niche. AML is known to alter stromal and vascular microenvironments, but these effects are difficult to separate from bulk leukemic burden and immune inflammation. Here, we use isogenic human AML xenografts with distinct LSC characteristics but comparable engraftment to define LSC-associated niche remodeling *in vivo*. Single-cell profiling revealed that LSC-high AML shifts the mesenchymal niche toward fibro-inflammatory states, expanding *Fmod+* fibroblasts and *Cd34*^+^ perivascular fibroblast-like cells while suppressing osteolineage differentiation. The leukemic compartment remained heterogeneous, with a specific MEP-like LSC population expressing niche-remodeling ligands including TGFB1, IL1B, and ANGPT1. LSC-high AML activated a TGFβ-responsive, CREB3L1-controlled fibroblast trajectory, and perturbing TGFβ signaling or CREB3L1 activation reduced stromal support for AML cells. These findings identify a specific LSC subtype as a source of niche-remodeling cues that shape specialized leukemia-supportive niches.

## INTRODUCTION

Acute myeloid leukemia (AML) is a heterogeneous blood malignancy characterized by the expansion of abnormal myeloid blasts in the bone marrow (BM) and peripheral blood. Despite advances in understanding its molecular and genetic drivers, AML remains a therapeutic challenge, with high relapse rates partly driven by leukemic stem cells (LSCs), a subset of therapy-resistant cells^1^. Due to the clonal expansion of leukemic cells, the BM niche is physically remodeled. Beyond this, recent evidence suggests that AML cells also actively reprogram the BM niche into a protective microenvironment that supports their survival and impairs normal hematopoiesis^2^, making it a potential target for therapeutic strategies.

The BM niche is a highly plastic system composed of mesenchymal stromal cells (MSCs), various specialized stromal cell types, endothelial cells, and non-cellular elements such as extracellular matrix (ECM), which collectively regulate hematopoiesis. The BM responds to internal and external stimuli through changes in cell states and cellular composition. AML contributes to this remodeling: for example, the secretion of TNF can lead to endosteal blood vessel loss and endothelial dysfunction^3,4^, and AML alters MSC differentiation, compromising their ability to support normal hematopoiesis^1,5,6^. Single-cell and spatial profiling studies have begun to map the complex interactions between the niche and hematopoietic stem cells (HSCs) and describe the functional heterogeneity of the stromal cells^5,7–11^. These stromal populations are fragile and highly sensitive, such that the cellular composition recovered is strongly affected by the isolation strategy applied. How LSCs specifically, rather than bulk leukemic cells or the underlying genetic profile, drive these niche alterations has remained difficult to resolve, as LSC content, mutational background, and disease burden are confounded in patient samples.

To address this gap, we used patient-derived xenograft (PDX) AML models with distinct LSC characteristics to directly determine the extent and nature of BM niche remodeling by AML LSCs. The model builds on our identification of hepatic leukemia factor (HLF) as a key regulator of LSCs in AML harboring mutations in *NPM1* (Nucleophosmin 1), *FLT3-ITD* (FMS-like tyrosine kinase 3), and *DNMT3A* (DNA methyltransferase 3a)^12^. Additionally, adhesion receptor GPR56 marks AML cells with high *in vivo* repopulating capacity irrespective of CD34 status^13^. Within the GPR56^+^ AML, two LSC compartments coexist and interconvert: a slowly cycling HLF-high CD34^+^GPR56^+^ population with a high LSC frequency, and a more rapidly cycling CD34^-^GPR56^+^ population of lower but still leukemia-initiating potential^12,14^, sustained by GPR56-driven co-activation of reciprocally antagonistic Hedgehog/TGFβ and Wnt programs^14^.

Our models include an isogenic pair of AML conditions differing in HLF: an HLF-high AML with slowly cycling CD34^+^GPR56^+^ LSCs (LSC-high) and a CRISPR/Cas9 HLF-knockout, which is characterized by a depletion of the CD34^+^GPR56^+^ population and their LSC balance shifts toward the faster-cycling, less-potent CD34^-^GPR56^+^ state (LSC-low). As LSCs cannot be maintained independently of their downstream progeny, both conditions comprise the full leukemic hierarchy and differ in LSC frequency.

Coupling single-cell RNA-seq, gene regulatory network inference, and imaging in these xenograft models and human primary MSC-AML co-cultures, we mapped niche remodeling at cellular and molecular resolution. We find that the presence of the slowly cycling, HLF-wild type LSC compartment rewires the stromal architecture towards a fibro-inflammatory state through a TGFβ-responsive transcriptional program. Cell-cell communication analysis identified TGFβ1 as the dominant LSC-derived cue, acting not on one niche cell type but convergently across both stroma and vasculature.

## RESULTS

### A defined isogenic PDX model and single-cell reference map of the AML bone marrow niche

Dissecting how leukemic stem cells (LSCs) remodel the bone marrow (BM) stromal niche is challenging due to the genetic diversity of AML, the poor correlation between mutational burden and functional LSC frequency^15^. To overcome these limitations, we leveraged a defined, isogenic patient-derived xenograft (PDX) model of AML with defined LSC frequencies. We xenotransplanted two matched AML conditions in immunodeficient NSGW41 mice^16,17^, which support multilineage human hematopoietic engraftment without irradiation-induced niche damage^18,19^: (i) an HLF-wild type AML with a high frequency of slowly cycling CD34^+^GPR56^+^ LSCs (LSC-high) and 2) a CRISPR/Cas9-mediated HLF knockout derivative containing mostly rapidly cycling CD34^-^GPR56^+^ LSCs (LSC-low). As additional references, we transplanted healthy cord blood (CB)-derived CD34^+^ HSPCs and included non-transplanted controls (**Fig. 1A**).

**Figure 1:**
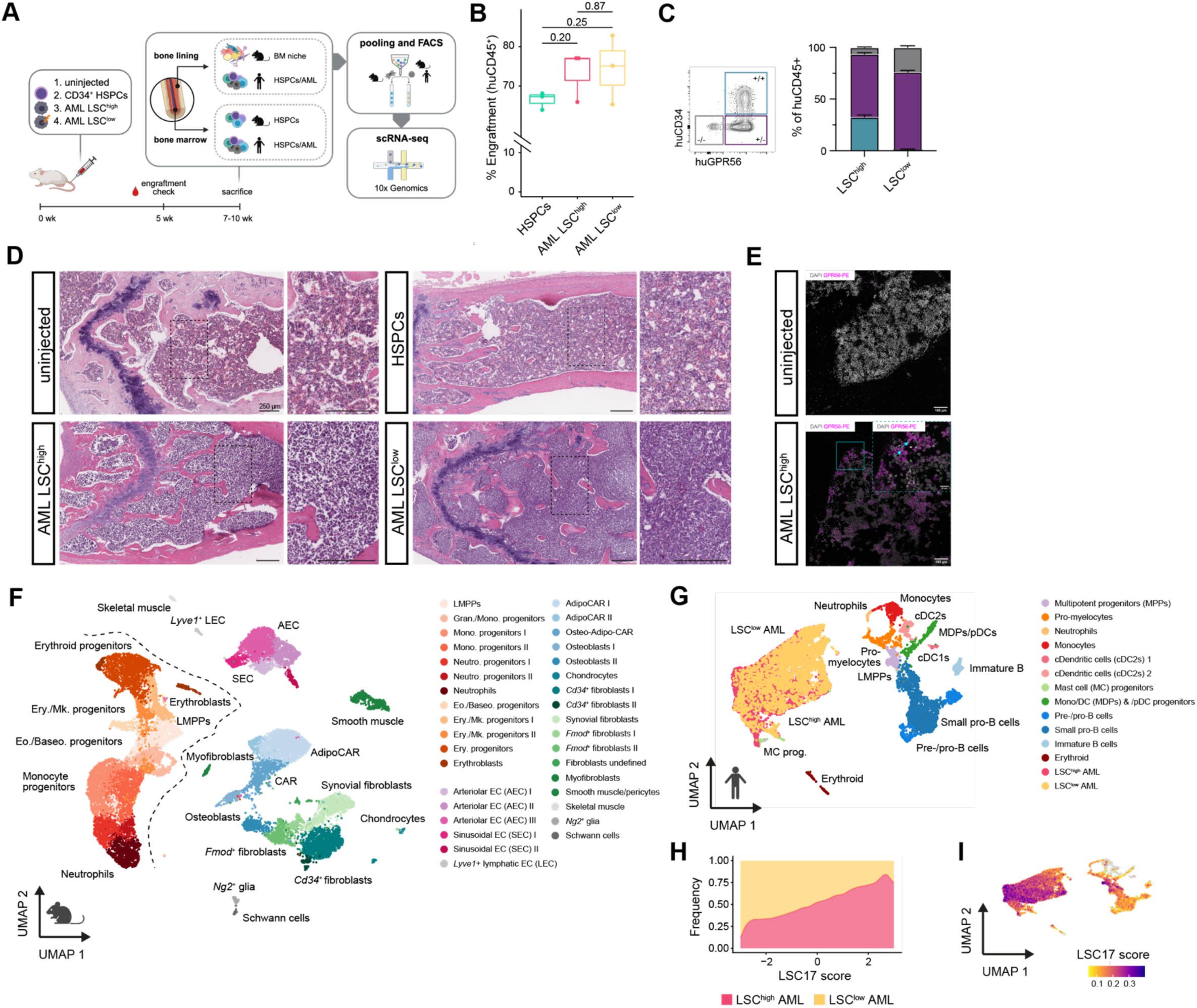
An isogenic PDX model with differential LSC burden enables integrated profiling of AML-driven bone marrow niche remodeling. (A) Experimental overview of the isogenic AML POX model. NSGW41 mice were transplanted with healthy cord blood (CB) CD34+ HSPCs, LSC-high AML, or LSC-low AML (isogenic HLF knockout), alongside non-transplanted controls. At sacrifice, bone-marrow and bone-lining compartments were isolated separately, FACS-purified, pooled (per condition), and processed for droplet-based scRNA-seq (1Ox Genomics, n = 3 mice/condition). (B) Flow cytometric quantification of human engraftment levels(% hCD45+) across transplanted conditions at 8 weeks post-transplantation. Engraftment was comparable across all conditions (n.s.; pairwise p-values indicated), confirming that downstream differences reflect LSC burden rather than graft size. (C) Flow cytometric quantification of the human AML conditions (% of hCD45+), showing the relative abundance of CD34+GPR56+ and CD34-GPR56+ LSCs and CD34-GPR56-blasts. Representative gating shown (left). (D) Representative H&E-stained sections of femoral BM FFPE from non-transplanted, CD34+ HSPCs, LSC-high AML, and LSC-low AML-transplanted mice. Boxed regions are shown at higher magnification (right panels). Scale bars: 250 µmin overview and insets. (E) Representative immunofluorescence staining of BM cryosections showing human AML infiltration, stained for hGPR56 (magenta) and nuclei (DAPI). Max projections are visualized. Scale bars: 100 and 20 µm. (F-G) UMAP visualization of integrated single-cell transcriptomes from mouse (F) and human (G) BM compartments, colored by annotated cell types. (H) Distribution of LSC17 gene signature score per cell stratified by LSC-high and LSC-low AML conditions. (I) UMAP of human cells colored by the LSC17 score.

We performed droplet-based scRNA-seq on both the hematopoietic and non-hematopoietic compartments 7 weeks post-transplantation, a time point at which overall engraftment levels reached 70-80% across all conditions, with no significant difference between LSC-high and LSC-low AML (**Fig. 1B**). Flow cytometric profiling confirmed a significantly expanded slowly-cycling CD34^+^GPR56^+^ LSC population in LSC-high AML relative to LSC-low AML (**Fig. 1B,C**, **SFig. 1A,B**). Moreover, western blot confirmed HLF KO in the LSC-low condition (**SFig. 1C**), altogether confirming the robustness of our PDX model system. Histological analysis (H&E) of femoral BM sections showed increasing marrow cellularity and blast-like infiltration in AML LSC-high and LSC-low conditions relative to uninjected and HSPC-engrafted controls (**Fig. 1D**), and immunofluorescence staining confirmed the presence of hGPR56^+^ leukemic cells, which are absent in uninjected controls (**Fig. 1E**).

To comprehensively study the BM niche remodeling, we isolated both stromal (mLin^-^/mCD71^-^/hCD45^-^) and endothelial cells (mCD31^+^) from digested bone chips using fow cytometry (**Methods**) and FACS strategy (**SFig. 1D**). Additionally, we enriched human and mouse HSPCs (respectively hCD45^+^/hCD34^+^ and mCD45^+^/mc-Kit^+^) and human AML cells (hCD45^+^) (**SFig. 1D**). Cells from individual mice per condition were barcoded with sample-specific oligo-conjugated antibodies and pooled for droplet-based scRNA-seq.

After species– and sample-level demultiplexing, quality control, and filtering (**SFig. 2A,B**), we obtained 22,981 mouse cells and 14,316 human cells. Unsupervised clustering and annotation based on canonical marker genes and manual annotation identified distinct hematopoietic, endothelial, and stromal populations (**Fig. 1F,G, SFig. 3A-C**, {https://apps.embl.de/hlfshiny/; Username: authors; Password: hlfhlf}. In line with our flow cytometry data, LSC-high AML cells exhibited significantly elevated LSC17 gene signature^20^ scores compared to LSC-low AML (**Fig. 1H,I**; unpaired Student’s t-test; p.adj < 2e-16), validating the transcriptional distinction between the two leukemic states.

**Figure 2:**
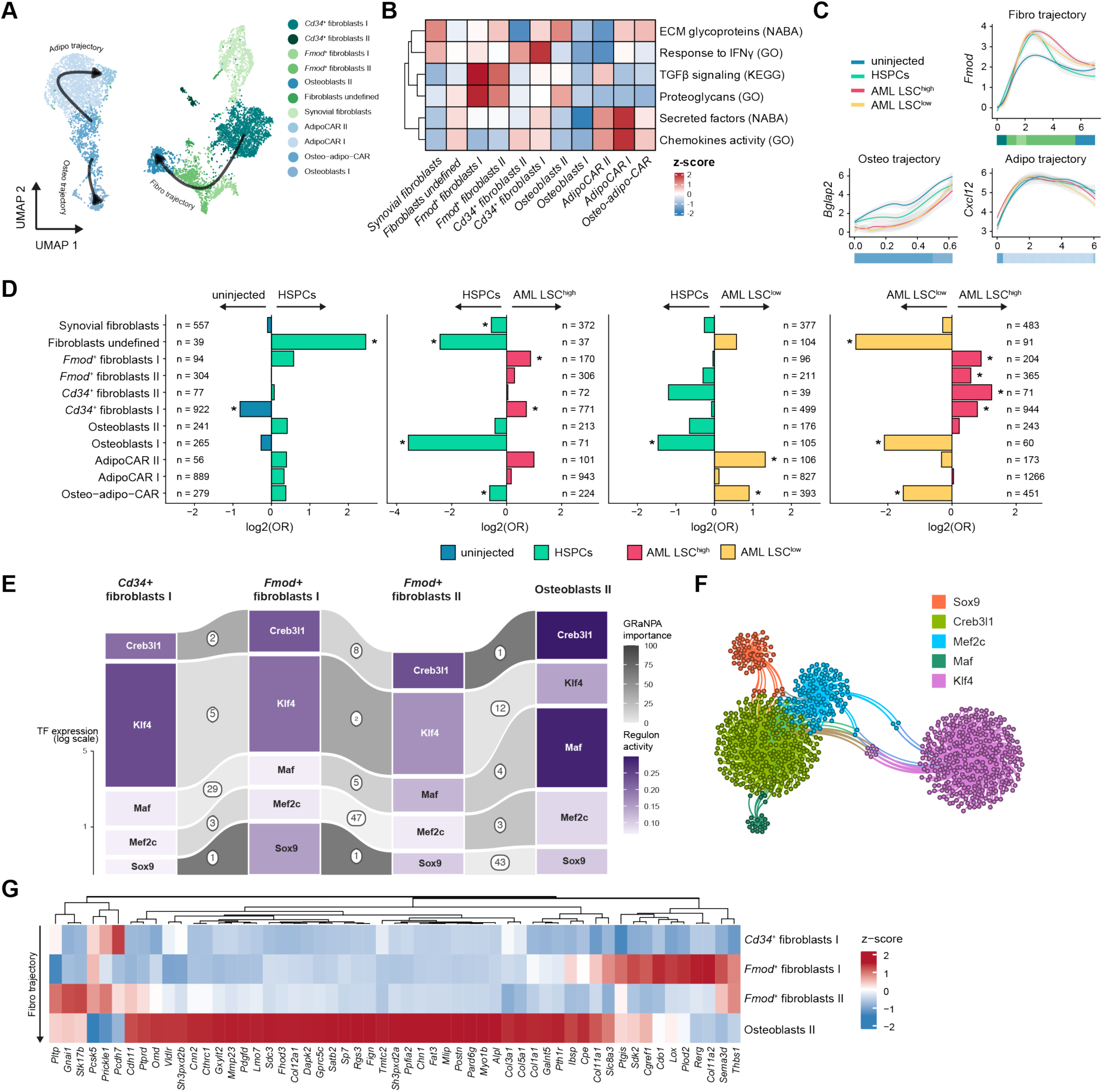
Intrinsic remodeling of the stromal compartment in LSC-high and LSC-low AML. (A) UMAP representation of stromal compartment, colored by annotated cell type. Black arrows indicate averaged pseudotime trajectories inferred with Monocle3 (see **SFig. 6A**). (B) Average expression score of selected gene signatures is shown for each stromal cell type as z-score across rows. Information regarding the origin of each gene-set is GO: Gene ontology; KEGG: Kyoto Encyclopaedia of Genes and Genomes; NABA: matrisome categories (Naba et al, 2012) (C) Normalized expression of selected lineage-specific marker genes is shown for the Osteo-*(Bglap2),* Fibro-*(Fmod),* and Adipo-trajectory *(Cxc/12)* markers, stratified by condition: uninjected (blue), and injected with HSPCs (green), LSC-high AML (red), LSC-low AML (yellow). (D) Remodeling of the stromal compartment between conditions is shown as odds ratios for: HSPC vs uninjected (left), LSC-low AML vs HSPC (second from left), LSC-high AML vs HSPC (second from right), LSC-high AML vs LSC-low AML (right). The analysis was performed per cluster using Fisher’s exact test, adjusted p-value after Bonferroni correction are shown(*: <0.05). (E) Transcription factor (TF) importance as quantified by GRaNPA (Kamal et al., 2023) for pairwise cell-state transitions along the Fibro-trajectory. For each TF, the box height encodes TF expression (log scale), box fill (purple) encodes regulon activity quantified as average expression of their inferred target genes (SCENIC(Aibar et al., 2017)). and the shade (grey scale) of the connecting ribbons depicts the GRaNPA-derived TF importance for that given transition. For the visualization, we included only TFs ranked top 5 by importance (actual rank shown as a circled number) in one of the transitions and appearing at least in 2 transitions. (F) Regulon of the most important selected TFs along the Fibro trajectory is displayed as a network of the TFs’ target genes. (G) Expression of genes in the Creb3I1 regulon is shown as per-gene z-scores for each cell state along the fibro trajectory.

**Figure 3:**
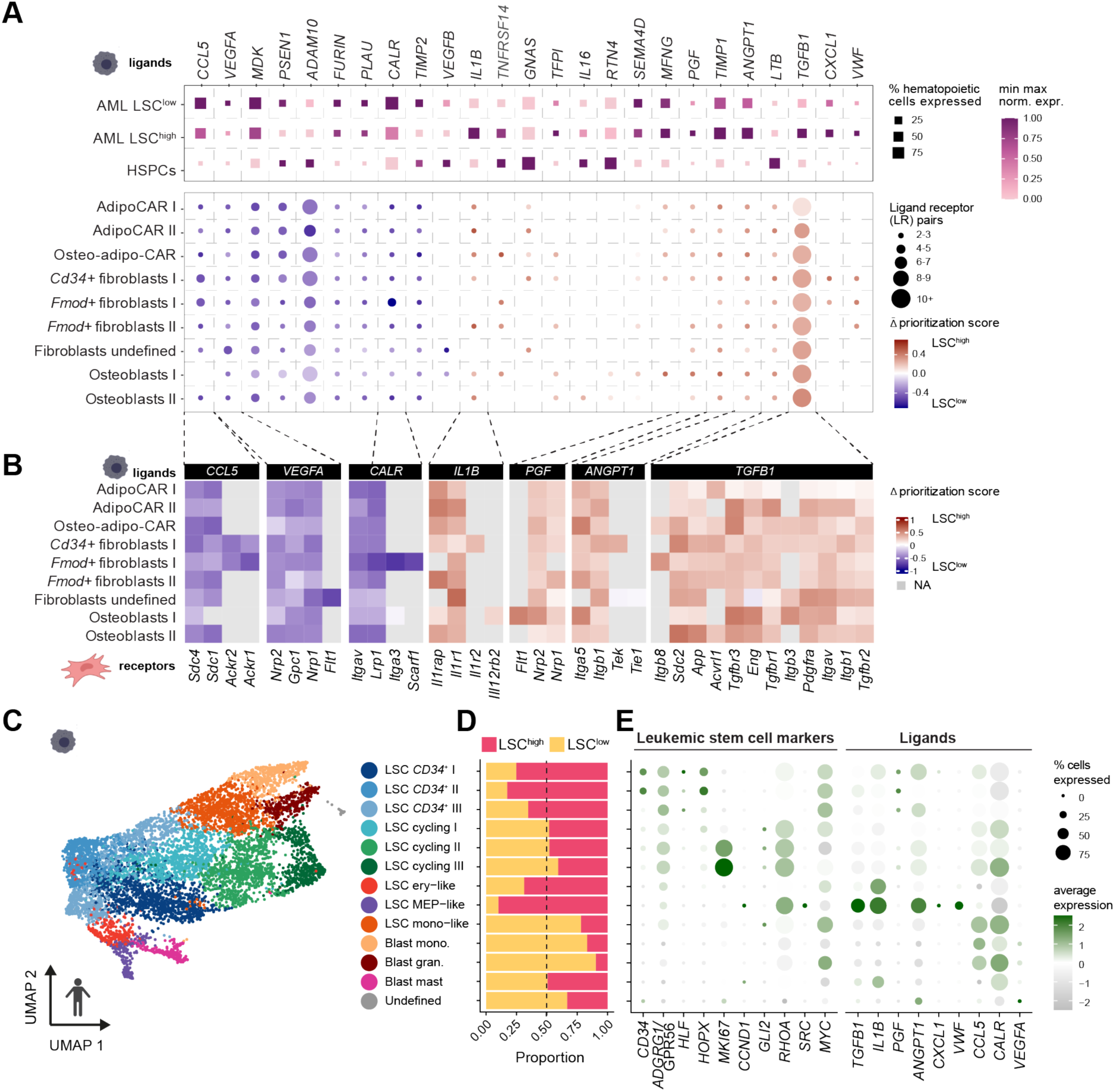
Extrinsic signals from AML to the stromal compartment. (A) Top: expression of ligands in AML LSC-low, LSC-high, and HSPCs, shown for the 25 ligands with the highest average differential prioritization score between LSC-high and LSC-low AML as quantified by MultiNicheNet(Browaeys, 2023) **(Methods).** Square size reflects the percentage of cells expressing the ligand, and color reflects the min-max normalized expression. Bottom: differential prioritization score for each ligand, averaged across all receptors, stratified by stromal cell types. Dot size reflects the number of ligand-receptor (LR) pairs, and color reflects the average differential prioritization score (red prioritized in LSC-high; blue: prioritized in LSC-low). (B) Differential prioritization scores for selected ligands across all their receptors, stratified by stromal cell types; colors as in A, bottom (C) UMAP representation of the AML cells colored by subcluster, annotated as distinct AML cell states (annotations in **SFig. 8**). (D) Fraction of cells originating from LSC-high (red) vs LSC-low (yellow) AML conditions for each AML subcluster. Dashed line at 50%. (E) Average expression of selected leukemic stem cell marker genes (left) and ligands (right) is shown for each AML subtype. Dot size reflects the percentage of cells expressing the genes, color represents the scaled average gene expression.

Within the mouse hematopoietic compartment, we resolved 12 clusters encompassing megakaryocytic progenitors and erythroblasts (Ery/Mk progenitors), neutrophil and monocyte progenitors, and eosino/basophil progenitors (Eo./Baso. progenitors) (**Fig. 1F, SFig. 3B**). Consistent with the NSGW41 genetic background, which impairs lymphoid maturation^17,21^, we identified a cluster of multipotent progenitors (LMPPs; *Kit*, *Msi2*, *Pim1*) but no further committed lymphoid progenitors. In contrast, the xenotransplanted human CD34^+^ HSPCs gave rise to both myeloid and lymphoid lineages, with the lymphoid output largely restricted to the B cell lineage (**Fig. 1G, SFig. 3C**) as previously shown for NSGW41 mice^16,17^.

In the non-hematopoietic compartment, we identified smooth muscle cells/pericytes (*Myh11, Mustn1, Tagln, Acta2*), Schwann cells (*Mal, Mag*), Ng2^+^ glial cells (*Ng2*/*Cspg4*, *Kcna1* positive), myofibroblasts (*Acta2, Myf5*), and multiple endothelial subtypes (*Cdh5, Pecam1*), including lymphatic (LEC; *Prox1, Lyve1*^22^), sinusoidal (SEC; *Emcn*), and arteriolar (AEC; *Ly6a*) endothelium (**Fig. 1F, SFig. 3A**). Within the mesenchymal compartment, we annotated 11 mesenchymal populations, including osteogenic clusters (Osteoblast I/II); *Bglap*, *Col1a1*), chondrocytes (*Sox9, Acan*), *Cxcl12*-abundant reticular (CAR) lineage subsets, and fibroblasts. Within the CAR lineage (*Cxcl12*, *Kitl, Lepr*), we resolved one Adipo-Osteo-CAR population (*Alpl, Mmp13*) and two Adipo-CAR subsets (Adipo-CARI/II; *Adipoq*). Fibroblasts segregated into two *Cd34^+^* fibroblasts (*Cd34*^+^ fibroblasts I/II), two *Fmod* fibroblasts (*Fmod*^+^ fibroblasts I/II), an undefined fibroblast cluster (Fibroblasts_undefined), as well as a distinct population of synovial-like fibroblasts (*Clic5, Prg4*), likely originating from joint-adjacent bone surfaces^23,24^. We also identified a cluster of skeletal muscle cells (*Ckm, Ttn*), corresponding to cells from the periosteal or outer bone surface, which were excluded from downstream analyses.

To contextualize the NSGW41 BM niche composition, we integrated our stromal and endothelial data with three published mouse BM stromal atlases on C57BL/6J mice^5,7,9^. This analysis demonstrated broad conservation of major niche populations across mouse strains (**SFig. 4A**), including SECs and AECs, perivascular cells, fibroblasts, and various CAR subsets, while also revealing dataset-specific biases driven by diverse enrichment strategies used (**SFig. 4B,C**). CAR subsets consistently spanned adipogenic, osteogenic, or mixed lineage programs, supporting a continuum rather than discrete lineage commitment states (**SFig. 4C**). By mapping the mouse stromal cell populations to a human reference^25^, we recapitulated the CAR, the osteogenic, and fibroblastic lineages, suggesting similar stromal populations exist in mouse and human (**SFig. 4D**).

**Figure 4:**
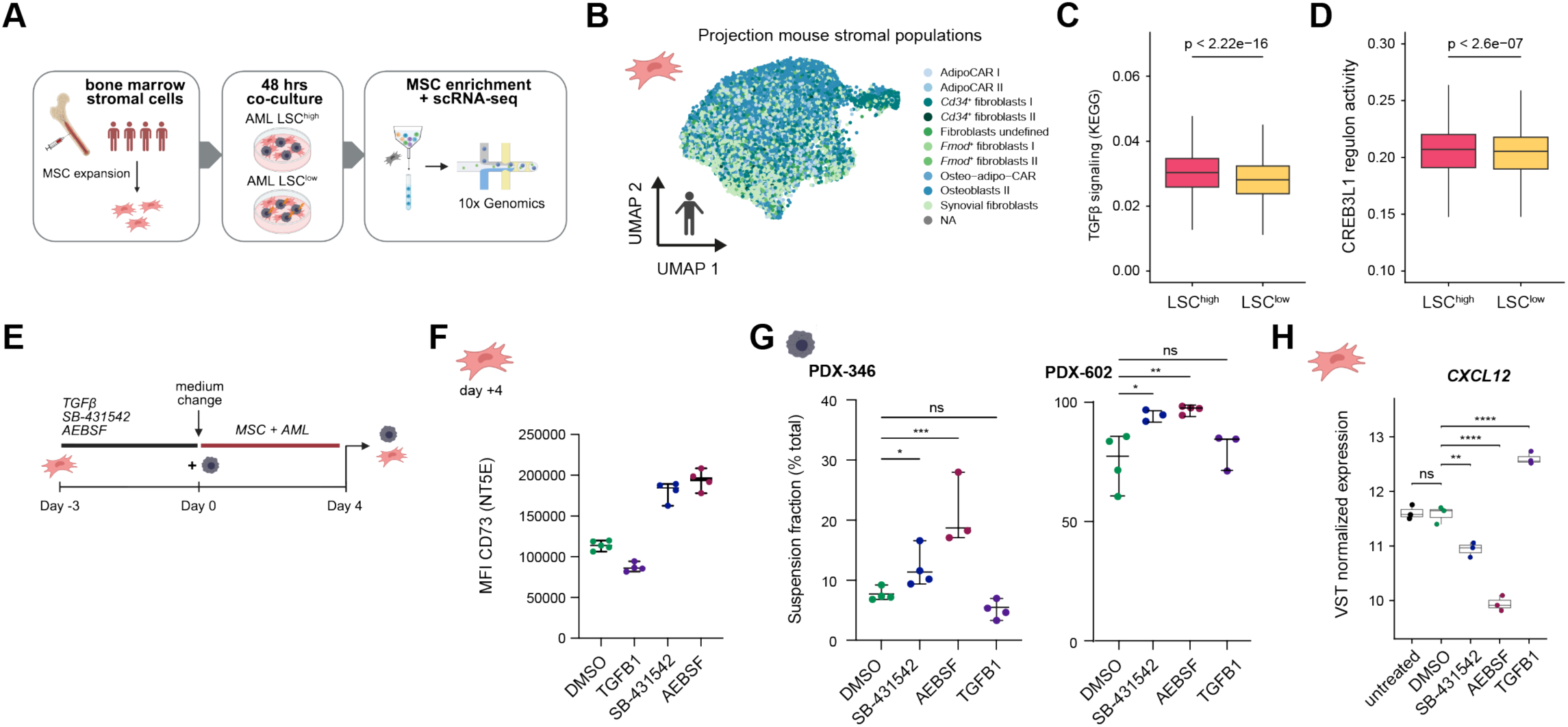
Modulation of TGFl3 and CREB3L1 impacts MSC-support function *in vitro*. (A) Schematic of the co-culture experiment: human primary BM MSCs from four independent donors were co-cultured for 48 hours with LSC-high or LSC-low AML cells, followed by MSC enrichment and scRNA-seq. (B) UMAP of scRNA-seq profiles from human MSCs recovered after co-culture, colored by projected identity based on the mouse stromal cell taxonomy defined in this study (Fig. 1). Human MSC subpopulations recapitulate the stromal diversity observed in the mouse xenograft model. (C,O) TGFl3pathway activity (C; TGFl3 signaling KEGG gene set score, as in **Fig. 2B**) and CREB3L1 regulon activity (O; regulon as defined in Fig. 2G) in human MSCs co-cultured with LSC-high (pink) versus LSC-low (yellow) AML. P-values from the Wilcoxon rank-sum test. (E) Schematic of the TGFl3perturbation co-culture experiment. Human primary BM MSCs from a healthy donor were pre-conditioned for 3 days with TGFl31 (10 ng/ml), the TGFl3type I receptor kinase inhibitor SB-431542 (10 µM), the CREB3L1 cleavage inhibitor AEBSF (250 µ **M),** or OMSO as a vehicle control. After medium change, AML cells were added and co-cultured for 4 days, after which flow cytometry was performed on both fractions. (F) Surface expression of C073 (NT5E) on MSCs measured by flow cytometry across the four pre-conditioning conditions at day 4 (n=4 replicates per condition). (G) Fraction of AML cells recovered in the suspension (non-adherent) fraction relative to total AML cells (adherent+ suspension), shown for two AML POX lines (POX-346, left; POX-602, right) across the 4 pre-conditioning conditions (n=4 replicates per condition per POX). One-way ANOVA followed by Benjamini, Krieger, and Yekutieli for multiple comparisons; q values are shown. *q<0.05, **q<0.01, ***q<0.001, ns = not significant. (H) VST-normalized CXCL12 expression in MSCs across the four pre-conditioning conditions and untreated control (n=3 donors, also **SFig. 7A**). Adjusted P values from a OESeq2 Wald test are shown. **p<0.01, ****p<0.0001, ns =not significant.

We further inferred spatial niche localization by mapping our single-cell profiles to a published bulk RNA-seq dataset of laser-capture microdissected (LCM) BM regions of C57BL/6J mice (**SFig. 4E**)^7^, encompassing the endosteal, sub-endosteal, sinusoidal, and arteriolar zones. Osteoblasts and osteo-CAR populations aligned with endosteal and sub-endosteal signatures, whereas the other populations rather matched uniformly across all compartments, with the perivascular CAR and smooth muscle populations preferentially in the vasculature compartments (**SFig. 4E**). Notably, the *Cd34^+^* fibroblasts I expressed *Pi16,* which had previously been associated with a high stemness potential in fibroblasts^26^ as well as perivascular fibroblasts in bone marrow and other tissues^7,27^, in line with the predicted location in the arteriolar compartment (**SFig. 4E**). Together, these analyses establish a comprehensive, spatially informed reference map of the NSGW41 BM niche, forming the foundation for subsequent analyses of AML-driven niche remodeling.

### Bone marrow stromal remodeling toward a pro-fibrotic state in LSC-high AML

To further characterize the individual stromal niche cells (**Fig. 2A)**, we quantified the expression of curated signatures with relevance to BM stromal function in each of the stromal cell populations. The *Fmod*^+^ fibroblast I/II populations were expressing a pro-fibrotic program, scoring highest for TGFβ signaling, ECM glycoproteins, and proteoglycans, whereas the *Cd34*^+^ fibroblasts were defined by an IFNγ-response and chemokine signature, indicative of an inflammatory, immunomodulatory identity (**Fig. 2B**). The CAR/Adipo-CAR populations were defined by secreted factor and chemokine programs. Projection of an HSPC-support signature (**SFig. 5A,B**, 46 genes, **Supplementary Table 1**,^28^) including key factors *Kitl*^29^*, Cxcl12, Il7*^30^*, Igf1*^31^*, Csf1*^32^, and *Bmp4*^33^ (**SFig. 5A**) confirmed that CAR populations held the strongest hematopoietic-support potential, whereas fibroblasts displayed variable supportive capacity depending on the subtype (**SFig. 5B**).

**Figure 5:**
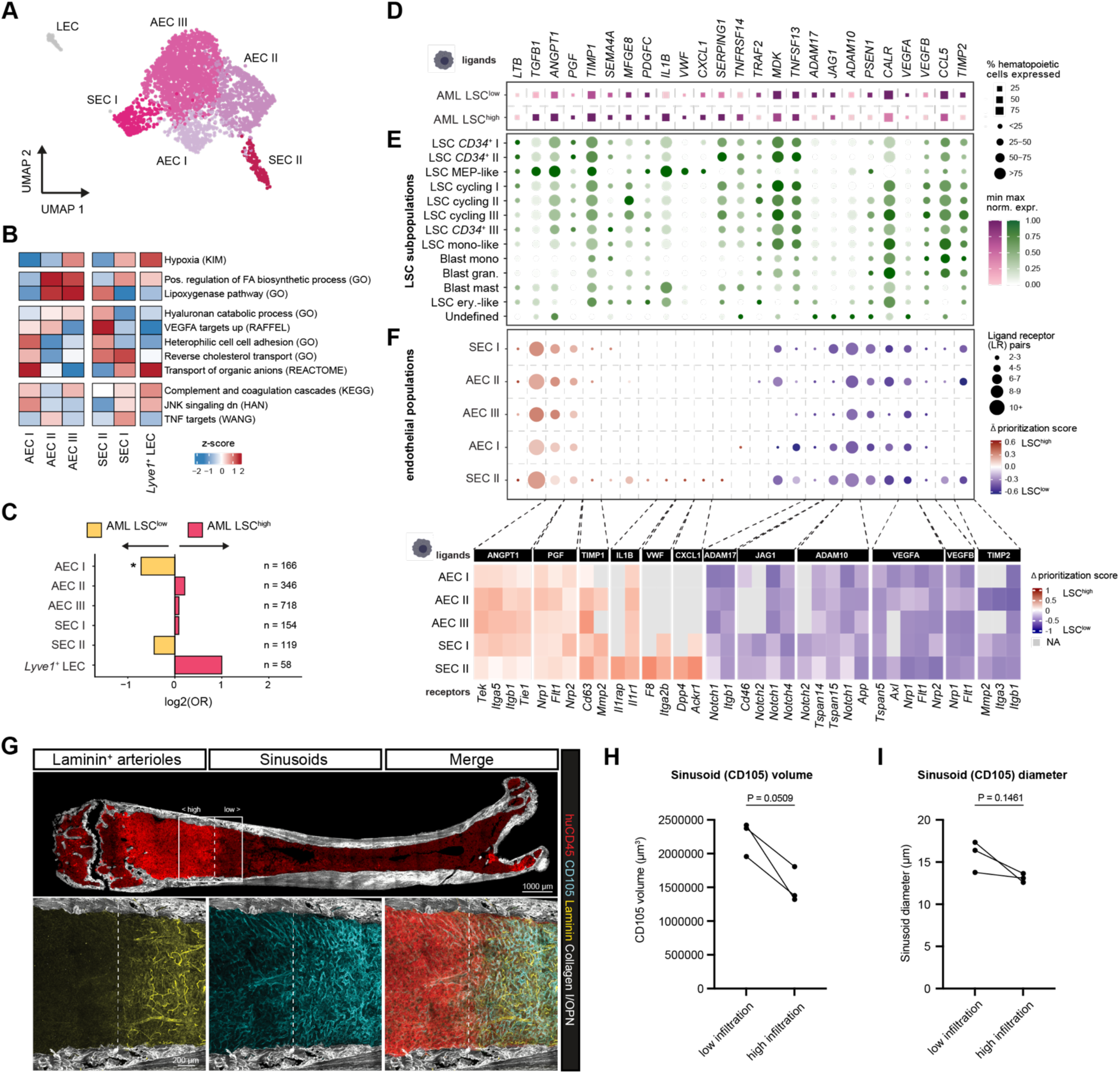
LSC-state– and infiltration-level dependent remodeling of the bone marrow vasculature network. (A) UMAP embedding of endothelial cells recovered from NSGW41 co-cultures across all conditions, annotated by subpopulation: arteriolar ECs (AEC 1-111), sinusoidal ECs (SEC 1-11), lymphatic-like ECs (LEC), and *Lyve1+* ECs. (B) Gene set enrichment heatmap showing selected differentially enriched pathways across endothelial subpopulations and conditions (LSC-high vs. LSC-low AML). Z-scores reflect normalized enrichment scores per cell population. (C) Odds ratio (OR) plot showing relative abundance of endothelial subpopulations in LSC-high (pink) and LSC-low (yellow) AML conditions compared to HSPC-injected controls. Bars represent log2(OR); asterisk indicates statistical significance (p < 0.05). Cell numbers (n) indicate total cells per subpopulation across conditions. (D) Dot plot showing expression of candidate MultiNicheNet ligands in AML LSC-low and LSC-high sender populations. Dot size reflects the percentage of hematopoietic cells expressing the ligand; color intensity reflects normalized mean expression. (E) Dot plot showing ligand expression across LSC subpopulations and blast populations identified within the LSC-high condition, for the same candidate ligands as in {D). (F) MultiNicheNet ligand-receptor analysis for endothelial receiver populations. Upper panel: dot plot showing the number of ligand-receptor pairs (dot size) and differential prioritization score (color: red = LSC-high enriched, blue= LSC-low enriched) for top-ranked ligands across endothelial subpopulations. Lower panel: heatmap of delta prioritization scores for selected ligands and their receptors across endothelial populations. (G) Representative image from volumetric (220 µm) tissue-wide fluorescence imaging of a femoral BM sample from a 7% engrafted mouse, stained for human CD45 (AML cells, red), CD105 (sinusoidal endothelium, cyan). Laminin (basement membrane/arterioles, yellow), and Collagen type 1/OPN (bone surfaces, white). Upper panel shows the full femoral section; lower panels show magnification of the boxed area, with a high-infiltration (left of dashed line) and low-infiltration (right of dashed line) region. Max projection is visualized. Scale bars: 1000 µm and 200 µm. (H) Sinusoidal volume (CD105+ volume, µm’) in low-versus high-infiltration regions, pooled across AML mice (n“3mice). Each data point represents the mean of all regions quantified per mouse per infiltration category. Lines connect paired measurements from the same mouse. P“0.0509, paired t-test. Full regional measurements are shown in **SFig. 100**. (I) Mean sinusoid diameter (µm) in low-versus high-infiltration regions, pooled across AML mice (n=3 mice). Each data point represents the mean of all regions (individual sinusoids) quantified per mouse per infiltration category. Lines connect paired measurements from the same mouse. P=0.1461, paired I-test. Individual sinusoid diameters for every region are shown in SFig. 10E.

Next, we characterized the lineage relationships of the BM stromal compartment, by inferring pseudotime using *Monocle3*^34^ (**SFig. 6A**). This identified three differentiation trajectories: two originated from CAR cells, bifurcating into adipogenic and osteogenic fates, and a fibrogenic trajectory starting from stem-like *Cd34*⁺ fibroblast I through *Fmod*^+^ fibroblasts I/II states, to a second osteogenic endpoint, Osteoblasts II (**Fig. 2A,C**). The two osteoblast endpoints, both *Bglap^+^* and *Col1a1^+^,* suggest that different osteoblast subsets may originate from distinct progenitor populations. This is consistent with recent work suggesting two sources of stromal progenitors in the mouse BM, both present in the bone lining fraction and able to give rise to convergent osteoblast populations through developmentally distinct differentiation trajectories^35^. The synovial fibroblasts formed a separate cluster with no direct lineage relationship to endosteal– and marrow-resident stroma. The expression of representative genes for each lineage (*Bglap2*; osteo-CAR trajectory*, Fmod*; fibroblast trajectory*, Cxcl12*; adipo trajectory) split by injected cell type suggested a loss of the CAR-lineage osteoblastic differentiation upon engraftment of AML versus healthy HSPCs, while the fibroblastic lineage marker *Fmod* was highest in mice injected with LSC-high AML cells (**Fig. 2C**).

To quantify the effect of engrafted cells on stromal remodeling, we determined relative population abundances using odds ratios (**Fig. 2D**). HSPC-engrafted controls differed only minimally from non-engrafted mice, limited to a small increase in smooth muscle cells and a small fibroblast subset (Fibroblasts undefined, n=39 cells), defining a near-homeostatic baseline. Against this baseline, LSC-high AML showed significant remodeling of the stromal compartment along the fibrogenic axis: both the pro-fibrotic *Fmod*^+^ fibroblasts I and inflammatory *Cd34*^+^ fibroblasts I were expanded, at the expense of Osteoblast I and fibroblasts-undefined. LSC-low AML also showed a reduction in the osteoblast I compartment, but, in contrast to LSC-high AML, showed an increase in AdipoCAR II rather than fibroblast subsets. A switch from osteo-to more adipogenic differentiation of the niche has been described for AML in general^5^. Direct comparison between LSC-high and –low AML confirmed the distinct effects of LSC-high and LSC-low AML on the niche: all *Cd34*^+^/*Fmod*^+^ fibroblast subsets were enriched in LSC-high AML, whereas the osteo-adipo-CARs were enriched in the LSC-low condition (**Fig. 2D**). Together, these data suggest that LSC-high AML drives the niche toward a fibro-inflammatory state, whereas LSC-low AML rather promotes an increased abundance of Adipo-CAR.

To identify the transcriptional regulators underlying the observed fibroblast fate transitions, particularly towards the more pro-fibrotic *Fmod*+ fibroblasts I/II populations, we inferred BM stroma-specific gene regulatory networks using *SCENIC*^36^ and used *GRaNPA* (Gene Regulatory Network Performance Analysis^37^), a machine learning tool, to identify the transcription factors (TFs) that were most predictive for the *Cd34*^+^ to *Fmod*^+^ to Osteoblasts II cell state transitions (**Fig. 2E**). Briefly, we quantified differential gene expression between consecutive pairs of cell states along the fibroblast trajectory, and used GRaNPA to assess how well the GRN predicts differential expression, and to identify the TFs that best explain expression changes for each transition. Overall, the GRN predicted differential expression with an R^2^ between 0.20-0.33 (**SFig. 6B, Methods**). Among the most predictive TFs were Creb3l1, Klf4, Maf, Mef2c, and Sox9 (**Fig. 2E**). Sox9 was the most important TF for the early transitions, with regulon activity highest at the *Fmod*^+^ fibroblast I state. Klf4 was also important early in the lineage, with regulon activity decreasing continuously, a pattern compatible with an anti-fibrotic role that has been suggested for KLF4 in lung fibroblasts^38^. MAF became important only at the final state transition, also reflected by its highest regulon activity in the Osteoblast II cell state. This is consistent with its established role in osteogenesis, cooperating with Runx2^39^. Mef2c, also previously implicated in osteoblast differentiation^40^, was important at the beginning and towards the last transition. In contrast, Creb3l1 was consistently among the top predictors across the fibrogenic trajectory, with a progressively increasing regulon activity. Its regulon was mainly non-overlapping with the other TFs, except for Mef2c (**Fig. 2F**), and was enriched for ECM, bone formation, fibroblasts, and fibrosis-like processes (**Fig. 2G, SFig. 6C**).

Creb3l1 showed a cell-type-dependent regulon activity along the trajectory, engaging distinct target genes in different stromal states. The majority of its targets reached their highest expression in the Osteoblast II state, but a distinct gene subset, including *Thbs1*, was specifically enriched in the *Fmod^+^*fibroblasts, the population expanded in LSC-high engrafted mice. Thbs1 promotes cleaving of latent TGFB1 in the ECM^41^ and has been found elevated in tumor microenvironments, including the bone marrow^28,42^.

To test this coupling of TGFB signaling, CREB3L1, and THBS1 directly, we treated primary human BM MSCs with TGFβ1, alone or together with the TGFBR1 inhibitor SB-431542, for 4 days, and profiled them by RNA-seq (**SFig. 7A-C**). TGFβ1 induced both *CREB3L1* and *THBS1*, and TGFβR1 inhibition attenuated this induction, placing both genes downstream of TGFβ signaling in human MSCs (**SFig. 7D**). Because THBS1 is itself an activator of latent TGFβ, these results point to a possible positive feedback loop to TGFβ in the ECM, engaging a CREB3L1/THBS1 program that liberates additional active TGFβ, progressively reinforcing the pro-fibrotic niche. This, in turn, raised the question of which leukemic signals initiate this circuit.

In summary, we found that LSC-high AML induced specific stromal remodeling compared with healthy HSPCs and LSC-low AML, resulting in an increased abundance of *Fmod*^+^ fibroblasts, which are characterized by high TGFβ-response driven partly through Creb3l1.

### Extrinsic drivers of stromal remodeling

To better understand how the LSC-high AML remodels the stromal niche, we next inferred cell-cell communication networks. To this end, we used MultiNicheNet^43,44^ to infer condition-specific communication between AML cells and stromal populations, converting human genes to mouse orthologs. MultiNicheNet prioritizes ligand-receptor interactions by integrating ligand expression in sender cells (here AML or HSPCs), receptor expression in receiver cells (here stromal cells), and the predicted ability of each ligand to explain downstream transcriptional responses in the receiver population, here defined as differentially expressed genes in LSC-high vs LSC-low AML conditions. The resulting prioritization score for LSC-high vs LSC-low identified TGFB1, IL1B, PGF, and ANGPT1 as the most active ligands in the LSC-high AML (**Fig. 3A**).

*TGFB1* was highly expressed in LSC-high AML (**Fig. 3A**), consistent with our previous finding that LSC-enriched CD34^+^GPR56^+^ cells express elevated *TGFB1*^14^. It was the only ligand that broadly interacted with all stromal populations via more than three receptors: Tgfbr1/2 receptor complexes, with Tgfbr3 and endoglin (Eng) acting as co-receptor and integrins (e.g., *Itgav, Itgb1*) as enhancers of the overall TGFβ signaling across stromal lineages (**Fig. 3B**). In particular, the predicted ligand-receptor pairs with TGFB1 are associated with activating Smad2/3-driven gene programs, which suppress adipogenesis and antagonize Smad1/5/8-driven gene programs downstream of BMP signaling that are associated with a more pro-adipogenic state^45^. The fibroblast populations expanded in LSC-high BM themselves showed increased TGFβ signaling activity (**Fig. 2B**), pointing to a positive feedback loop that reinforces TGFβ pathway activation within the leukemic microenvironment.

The inflammatory cytokine IL1B also showed higher predicted activity in LSC-high AML, signaling through Il1r1 across all stromal populations and additionally through Il1r2 in *Cd34^+^* fibroblasts I (**Fig. 3A-B**). Since Il1r2 functions as a decoy receptor that can sequester IL1 ligands rather than transducing a signal^46^, this suggests that *Cd34*^+^ fibroblasts I may not respond to but rather buffer IL1B-driven inflammation. In parallel, ANGPT1 showed broad predicted interactions with all stromal populations, mostly through integrin receptors. In *Cd34*^+^ fibroblasts I, ANGPT1 additionally signaled through its canonical receptor Tie2 (*Tek),* which points to a perivascular identity. This is consistent with their predicted arteriolar niche localization (**SFig. 4E**) and the known association of *Pi16*^+^ fibroblasts with perivascular and adventitial compartments^47–49^. Finally, PGF interacted broadly through *Nrp1/Nrp2* across stromal populations, while the PGF-Flt1 interactions were restricted to Osteoblasts I, indicating the VEGF-family signaling may preferentially target the osteolineage in the LSC-high niche.

In contrast, the ligands *CCL5*, *VEGFA*, and *CALR* showed higher predicted activity and expression in the LSC-low AML cells (**Fig. 3A**). CCL5 was predicted to interact with the stroma not through its canonical receptor Ccr5, but through syndecans (Sdc1/4) and the decoy chemokine receptors Ackr1/2, suggesting the stroma may scavenge or sequester CCL5 rather than mounting a direct inflammatory response (**Fig. 3B**). VEGFA was predicted to interact through the co-receptors Nrp1, Nrp2, and Gpc1 rather than the principal receptor Kdr (Vegfr2); as these co-receptors primarily bind VEGFA without transducing a canonical downstream signal, this points to stromal retention of VEGFA rather than active VEGF pathway activation. Finally, CALR showed higher predicted activity in LSC-low AML, interacting with stromal cells predominantly through integrins (Itgav, Itga3), consistent with a role in cell-matrix adhesion rather than the immunogenic cell death signaling associated with surface-presented CALR.

Together, these data suggest that LSCs serve as a dominant paracrine source of stroma-remodelling cues. Among these signals, TGFB1 emerges as a key mediator, driving a pro-fibrotic and differentiation-suppressive remodeling program in the niche and supporting an autocrine-paracrine TGFB circuit between LSCs and fibroblastic stroma that sustains a protective leukemic niche.

### Specific subpopulations of LSCs drive TGFβ-mediated niche remodeling

GPR56^+^ LSCs oscillate between a slowly cycling HLF-high CD34^+^GPR56^+^ population and a more rapidly cycling CD34^-^GPR56^+^ population^12,14^. The LSC-high AML model is therefore enriched for the slowly cycling LSC compartment, which is largely depleted due to HLF loss in the LSC-low condition^12^. To understand whether a specific LSC subpopulation, rather than the bulk AML compartment, was the source of the niche-remodeling ligands, we resolved the heterogeneity of the malignant compartments in the scRNA-seq data. We subclustered the AML compartment and assigned the populations into LSC and blast identities based on the LSC17 score^20^ together with canonical LSC markers of our AML model, including *ADGRG1* (*GPR56*), *CD34, HOPX*, and *HLF*, and blast identities assigned based on their top differentially expressed genes (**Fig. 3C-E, SFig. 8A**).

We identified nine LSC subpopulations and three blast-like populations. Among the LSCs, three non-cycling populations with high *CD34* expression (LSC *CD34^+^* I, II, III) showed low MKI67 expression and low S– and G2/M-phase scores (**Fig. 3D, SFig. 8B**), consistent with a slowly cycling or quiescent state. Three actively cycling populations (LSC cycling I, II, III) expressed high *MKI67* expression and have elevated S-score and G2/M-phase scores (**Methods**). A distinct MEP-like LSC population was characterized by a megakaryocytic-erythroid-like gene expression program, including *ITGB3*, *VWF*, *PF4*, *GATA1*, *G6B*, and *CD9,* alongside *GPR56* (**SFig. 8A**). Its MEP*-*like lineage bias is consistent with the CD34^-^GPR56^+^ LSC fraction that we previously identified as capable of reconstituting the CD34^+^ compartment under TGFβ stimulation and carrying the Hh/TGFβ effector signature (CCND1, SRC, RHOA, TGFB1)^14^(**Fig. 3E**). Notably, high *VWF* expression, one of the key MEP-genes in this cluster, was previously found enriched in GPR56-high AML and suppressed upon GPR56 knockdown^12,14^, further linking this population to the GPR56-high LSC compartment. Two additional LSC populations with moderate GPR56 and LSC17 scores were identified based on lineage-associated gene programs: mono-like LSCs (*TREM2, FCN1*), and ery-like LSCs (*GATA1, KLF1*). The remaining blast populations included monocytic (Blast mono.), granulocytic (Blast. gran.), and mast cell-like (*TPSAB1*, Blast mast) (**Fig. 3C-E, SFig. 8A,B**).

Because we separately enriched AML cells from bone-lining and marrow fractions and multiplexed samples by barcoded antibody hashing (**SFig. 8D**), we could assign subpopulations to each compartment (**SFig. 8E**). The LSC *CD34*^+^ I population was predominantly recovered from the lining fraction, whereas the remaining LSC and blast populations were more evenly distributed across the bone, consistent with prior observations that quiescent LSCs preferentially reside at the endosteal niche^50^.

Ligand expression analysis identified the MEP-like LSC I population as the dominant source of TGFβ, together with IL1B and ANGPT1, and was almost exclusively contributed by the LSC-high condition (**Fig. 3E**). This suggests a role for the HLF-dependent LSC hierarchy in maintaining this population (**Fig. 3D, SFig. 8C**). LSC *CD34*^+^ I/II/III and ery-like LSCs were significantly enriched in the LSC-high condition, whereas LSC-low blasts spanned a more diverse mix of monocytic and granulocytic (GMP-like) states (**Fig. 3D, SFig. 8C**). The ligands most active in low-LSC AML (*CCL5, CALR, and VEGFA)* were largely restricted to the blast populations enriched in LSC-low AML. Together, these findings link the dominant niche-remodeling signals to specific LSC subpopulations, rather than to the bulk leukemia, with one LSC subpopulation driving the *TGFB1*-dependent niche remodeling.

### Effect of LSC-high and LSC-low AML cells on human MSCs

To further corroborate the observed TGFB1-dependent niche remodeling in the human BM context, we assessed how *in vitro* cultured human BM MSCs were remodeled upon co-culture with LSC-high vs LSC-low AML. Specifically, we obtained MSCs from 4 donors and co-cultured them for 48 hrs with LSC-high or LSC-low AML, followed by scRNA-seq profiling (**Fig. 4A; SFig. 9A,B**). We recovered 11,555 cells with a mean of 1,556 cells per sample, per condition (**SFig. 9C**). Projecting the mouse stromal population identities onto the human MSCs recovered analogous populations (**Fig. 4B, SFig. 9D,E**). In line with the observation in the PDX models (**Fig. 3A**), the LSC-high-exposed MSCs were significantly enriched for *Fmod+* fibroblasts II (**SFig. 9D**) and had higher TGFβ signaling (**Fig. 4C**, p < 2.2 x 10^-16^), and CREB3L1-regulon activity (**Fig. 4D**, p < 2.6 x 10^-7^) than LSC-low-exposed MSCs. This provides an orthogonal line of evidence of the TGFβ-mediated niche-remodeling of the LSC-high AML.

We next asked whether TGFβ signaling and its downstream effector CREB3L1 play a causal role in the remodeling that promotes LSC maintenance (**Fig. 4E**). To directly test the functional role of TGFβ signaling in stromal support of AML, we pre-conditioned human primary BM MSCs with recombinant TGFβ1 or a pharmacological inhibitor. The TGFβ type I receptor kinase inhibitor SB-431542 prevents SMAD2/3 phosphorylation downstream of ALK4 and –5. To test the functional role of CREB3L1, we used AEBSF, a serine protease inhibitor that blocks the intramembrane proteolytic cleavage of CREB3L1, thereby preventing its activation and nuclear translocation^51^. MSCs were treated for up to 72 hrs with TGFβ1 or the inhibitors before co-culture. This pre-conditioning strategy was designed to simulate a remodeled stromal niche and to isolate stromal-intrinsic effects of TGFβ activation or blockade on subsequent AML-stroma interactions.

As expected, pathway perturbation modulated canonical TGFβ response genes in MSCs (**SFig. 7C**). Inhibition of TGFβ pathway activity using SB-431542 or AEBSF reduced expression of niche-support factors such as *CXCL12*, while surface CD73 (NT5E) was increased by both inhibitors and reduced by exogenous TGFβ treatment in AML co-cultures. As CD73 expression rises during osteogenic priming of BM MSCs, this is in line with previous studies that TGFβ shifts MSCs away from an osteogenic fate towards a more pro-fibrotic program (**Fig. 4B**)^52,53^. Morphological changes were also apparent, with MSCs adopting a thinner spindle-shaped morphology by the inhibitors (**SFig. 7B**). We next assessed how these stromal changes affected AML maintenance. Pre-conditioned MSCs were co-cultured with AML cells from two independent PDX AML models (PDX-346 and PDX-602), representing distinct patient-derived AML samples^54^. We quantified the AML fraction in both the non-adherent (suspension) and MSC-adherent compartments of PDX-346 and PDX-602 (**Fig. 4C**). Inhibition of TGFβ signaling (SB-431542) or CREB3L1 activation (AEBSF) significantly increased the non-adherent suspension fraction (**Fig. 4C**), indicating that stromal TGFβ signaling and CREB3L1 activity reduced adhesion of the AML cells to the MSC stromal layer. By contrast, exogenous TGFβ1 pre-conditioning produced only minimal additional increases in AML fraction, likely because the pathway was already saturated by endogenous AML-secreted TGFB1 during co-culture.

Together, these results support a model in which LSCs remodel the stromal microenvironment through a TGFβ-CREB3L1-dependent axis, creating a fibrotic, quiescent niche that sustains LSCs. Blocking TGFβ signaling, either at the SMAD2/3 phosphorylation step (SB-431542) or by preventing CREB3L1 activation (AEBSF), diminishes stromal support for LSCs, highlighting a positive feedback loop in which AML-derived TGFB1 reinforces fibroblast TGFβ activation (**Fig. 4D**).

### Remodeling of the vascular niche by AML and linked to LSC burden

Having identified LSC-associated remodeling of fibroblastic stromal populations, we next asked whether this remodeling extends to the vasculature. The BM vasculature is known to undergo structural and functional alterations in AML, including changes in sinusoidal and arteriolar endothelial cells and increased vascular permeability^3,4^. We therefore assessed vascular remodeling in relation to LSC burden in our PDX model using the single-cell dataset and complementary imaging of the femoral BM vasculature (**Fig. 5A**).

We identified six endothelial types: three arteriolar (AEC I-III), two sinusoidal (SEC I-II), and one lymphatic (LEC) population (**Fig. 5A**). The AEC states varied mainly in hypoxia-associated signatures (highest in AEC III), fatty-acid biosynthesis and lipid-related pathways (highest in AEC II/III), organic anion transport (highest in I), and heterophilic cell-cell adhesion (highest in AEC I). The SEC states differed in their VEGFA target activity (highest in II), heterophilic cell-cell interaction (highest in II), and hypoxia-associated programs (highest in I) (**Fig. 5B**).

Endothelial remodeling was less pronounced at the level of cellular composition than what we observed in the stroma. The main abundance difference was a relative enrichment of AEC I in LSC-low AML compared to LSC-high AML (**Fig. 5C**). Transplantation of HSPCs alone had minimal effects on the endothelial compartment, except for a modest reduction in the AEC I subset (**SFig. 10A**). Notably, in both AML conditions SEC populations show a consistent trend toward relative enrichment compared to HSPC-injected controls (**SFig. 10A**), which is a pattern consistent with increased sinusoidal microvessel density in AML patient samples^55–57^. However, gene set enrichment analysis on differentially expressed genes revealed distinct functional programs between endothelial cells exposed to LSC-high versus LSC-low AML, most prominently within SEC II (**Supplementary Table 2**). LSC-low AML induced programs of microtubule dynamics, spindle assembly, and cell cycle progression, indicating a more proliferative endothelial state. LSC-high AML instead induced chemokine-signaling programs together with gene sets related to contractile functions and cytoskeletal remodeling (**Supplementary Table 2**). Beyond SEC II, condition-dependent effects were limited to decreased collagen formation in AEC III and immune-response terms in LEC (**Supplementary Table 2**).

To identify the cues underlying these gene programs, we performed cell-cell communication analysis with MultiNicheNet between AML cells and EC subsets (**Fig. 5D-F**). Consistent with the chemokine gene programs induced, LSC-high AML engaged with SEC II predominantly through chemokine– and cytokine-mediated interactions, including inflammatory mediators *IL1B*, CXCL1, and *LTB*, *VWF*, *TNFRSF13* (**Fig. 5F**), and TGFB1 as the dominant ligand. LSC-low AML, in contrast, signaled to the endothelium through *Notch* receptors on endothelial cells, such as *MDK (Notch2), ADAM17 (Notch1), JAG1 (Notch1/2/4), ADAM10 (Notch1/2),* and *PSEN1 (Notch1),* alongside *VEGFA/VEGFB* and *CCL5. This is* in line with the role of Notch– and VEGF-induced signaling on endothelial homeostasis^58^. The ligands signaling from LSC-high AML were mostly expressed in the LSC-high specific *CD34+* I/II and LSC I populations (**Fig. 5E**), whereas LSC-low VEGF, MDK, and CCL5 were expressed more broadly across various blast and proliferative LSC (III/IV) compartments.

Finally, we examined the BM vasculature network *in situ* by 3D tissue-wide femoral imaging of an independent mouse cohort, stained for human CD45 (AML cells), CD105 (sinusoidal endothelium), Laminin (basement membrane), and Collagen type I and Osteopontin (bone surfaces) (**Fig. 5G, SFig. 10B**). In both LSC-high and LSC-low conditions, the vasculature was disrupted, showing altered vascular integrity (**SFig. 10CC**). Sinusoids were less connected and less voluminous in highly infiltrated regions compared to low infiltrated regions, as determined by their cumulative volume and the mean sinusoid diameter (**Fig. 5H,I, SFig. 10D,E**). Notably, the extent of sinusoidal vessel disruption correlated with leukemic burden, being most pronounced in regions with high AML infiltration (**Fig. 5G**). The Laminin^+^ BM arterioles were also reduced and discontinuous in highly infiltrated regions. Importantly, these changes were spatially restricted to regions with high AML density: even in highly engrafted femurs, areas of lower infiltration retain an apparent intact vascular architecture (**SFig. 10C**), suggesting that vascular disruption is at least partly a consequence of mechanical displacement from dense leukemic infiltration. This suggests that the endothelial impact of AML is highly spatially localized and that the disruption of the architecture may be due to highly proliferating LSC populations. Accordingly, the endothelial populations we recovered in our single-cell data likely derived predominantly from less-infiltrated regions, accounting for the comparatively subtle transcriptional changes.

Collectively, these data suggest that vascular remodeling in AML is shaped not necessarily by LSC burden but by local leukemic density. While LSC burden was associated with endothelial lineage changes and altered signaling cues, the spatially restricted disruption of vasculature indicates that local AML infiltration further contributes to this remodeling.

## DISCUSSION

Previous work has established that AML actively remodels the bone marrow microenvironment, including stromal, osteolineage, and endothelial niches^3–5,59,60^. Yet, the drivers of this remodeling remain difficult to resolve in patient samples, where genetic heterogeneity, variable leukemic burden, immune inflammation, and differences in LSC frequency are inherently confounded. Here, we used an isogenic AML xenograft model in which the leukemic conditions differ only in LSC burden^12^ while showing comparable overall engraftment. Because both conditions retain the full leukemic hierarchy, single-cell resolution was essential to move beyond a population-level association and localize niche-remodeling signals to specific LSC subpopulations rather than the bulk LSC or blast burden. This controlled setting allowed us to directly attribute niche remodeling to LSC state, and to dissect how LSC-high AML remodels the stromal and endothelial bone marrow niche.

An important feature of our model (NSGW41) is the absence of T cells^17^. This is relevant because recent studies in AML, MDS, and MPN have reported inflammatory BM niche states and suggested that T-cell-stromal interactions contribute to inflammatory niche remodeling(Chen et al., 2025; Prummel et al., 2025). Our findings in the NSGW41 model show that LSC-high AML is sufficient to induce a fibro-inflammatory stromal program even in the absence of adaptive immune cells, indicating that LSCs can directly impose fibrotic and inflammatory features onto the BM microenvironment. Specifically, we found that LSC-high AML led to an increased abundance of *Fmod^+^* fibroblasts and *Cd34^+^* fibroblasts with perivascular features, and reduced osteolineage differentiation. This is consistent with studies showing that myeloid malignancies can alter the balance between hematopoietic-supportive, osteogenic, and fibrotic stromal programs^5,61^. Thus, while overt bone marrow fibrosis is most prominently associated with MPN, fibrosis-like stromal activation, extracellular matrix remodeling, and loss of normal niche-support functions are increasingly recognized as features of AML and MDS-associated niche dysfunction.

Notably, HSPC engraftment alone induced only minimal stromal remodeling relative to non-transplanted controls in our model. The near-homeostatic HSPC-only baseline should be also interpreted with some caution. A recent study on the bone marrow niche composition upon human HSPC engraftment in the same NSGW41 model reported a shift of *Pdgfra*^+^ MSCs toward a *Lepr*^+^ adipo-primed state upon long-term HSPC engraftment (>20 weeks) within the *Cd51^+^Pdgfra^+^Sca1^-^*stromal fraction^18^, suggesting that HSPC-induced niche remodeling in this model may be time– and enrichment strategy-dependent, and that our baseline reflects and earlier stage of this process

Mechanistically, our data identified an LSC-mediated TGFβ–CREB3L1 axis as a potential driver of this fibroblast remodeling. TGFB1 was highly expressed in LSC-high AML and was predicted to signal broadly across stromal cell types, consistent with the induction of pro-fibrotic gene programs. The strongest TGFB1-expressing LSC subpopulation displayed an MEP-like gene expression program, including *VWF, PF4, G6B,* and *ITGA2B*. Similar megakaryocyte-erythroid-biased HSPCs have been reported in myelofibrosis, where they are associated with TGFβ-driven fibrotic remodeling in the BM^62–64^. Moreover, megakaryocyte-derived PF4 has been shown to reprogram the Gli1^+^ fibrosis-driving stromal cells in MPN^65^. This raises the possibility that an MEP-primed LSC state drives TGFB-dependent fibrotic niche remodeling also in AML.

In the stromal compartment, TGFβ–induced CREB3L1 emerged as a key regulator along the fibroblast trajectory, with target genes linked to extracellular matrix organization and fibroblast activation, consistent with previous work showing that CREB3L1 was required for TGFβ-driven collagen ECM gene activation^66^. This mechanism may be reinforced by TGFβ being commonly stored in the ECM as an inactive latent complex, with local activation seen as a critical step controlling pathway activity^67^. Thus, LSC-derived TGFB1 may act as an initiating signal that induces a CREB3L1-dependent fibroblast and ECM program, rather than as a soluble ligand acting in isolation. The induction of THBS1 (Thrombospondin-1) is particularly relevant because thrombospondin-1 can activate latent TGFβ, thereby increasing local TGFβ bioavailability, and was recently implicated in fibrotic remodeling in MPN^65^. Thus, the CREB3L1/THBS1 program identified in *Fmod*^+^ fibroblasts may create a feed-forward circuit in which LSC-derived TGFB1 induces a stromal matrix state that further amplifies local TGFβ activation and stabilizes a pro-fibrotic, LSC-supportive niche. Consistent with this model, perturbation of TGFβ signaling or CREB3L1 activation reduced stromal support for the CD34^+^GPR56^+^ LSC fraction *in vitro*.

The vascular niche showed a related but less pronounced remodeling on the gene expression level. Consistent with reports that AML disrupts bone marrow vasculature and endothelial function^3,4^, both AML conditions showed structural disruption with reduced sinusoidal integrity and discontinuous arteriolar structures, particularly in regions of high AML infiltration. This architectural disruption suggests local leukemic density and dependence on mechanical displacement of the vascular network. In parallel, transcriptional profiling of recovered endothelial cells showed condition-specific functional programs: LSC-high AML was associated with inflammatory, chemokine, cytoskeletal and contractility-associated programs, consistent with actomyosin-driven junctional remodeling and increased vascular permeability^3,4^. In contrast, LSC-low AML showed more proliferative and VEGF/Notch-linked endothelial programs. Thus, AML may affect the vasculature through two partly separable mechanisms: local architectural disruption associated with leukemic infiltration, and LSC-state-dependent signaling to endothelial cells, potentially driven by TGFβ and inflammatory ligands identified in our NicheNet analysis.

Apart from the niche remodeling, our observations also extend previous work on LSC plasticity. In a previous study, we showed that NPM1c/DNMT3A/FLT3-ITD AML contains distinct LSC compartments^12,14^, including an immature, slowly cycling CD34^+^GPR56^+^ fraction, and proposed that reciprocal transitions between LSC states sustain the LSC pool. Here, we recovered this heterogeneity *in vivo* on the single-cell level and revealed that certain programs that we had associated with high GPR56 and HLF expression are derived from very specific LSC subsets. Importantly, the MEP-like LSC population does not map cleanly onto either compartment previously defined by bulk RNA-seq and flow cytometry profiling: while its high TGFB1 and SRC expression is consistent with the CD34^+^GPR56^+^ signature, the absence of *CD34* more closely resembles the faster cycling CD34^−^GPR56^+^ compartment. This indicates that the CD34^-^GPR56^+^ population harbors various LSC subpopulations. Moreover, these LSC populations differed not only in cell-cycle state and stemness-associated programs, but also in their expression of niche-remodeling ligands, including TGFB1. Our study positions HLF as a key driver of this TGFB1-expressing MEP-like LSC population. Since HLF itself is not detected within this population, this suggests that HLF is required upstream to establish this LSC state. This HLF-dependency may also be relevant for other HLF-driven entities such as TCF3::HLF childhood ALL, which has a dismal prognosis and has been shown to have an immature HLF-associated HSC-like gene program^68,69^.

LSCs are recognized as a major source of therapy resistance and relapse, in part through inherent resistance mechanisms including phenotype plasticity and dormancy^70^. Recent work has further defined functionally distinct LSC subtypes in primary AML patient samples with different sensitivity to venetoclax/azacitidine, among which a MEP-LSC subtype emerged under therapeutic pressure^71^. Our data raise the complementary possibility that LSC subtypes may additionally differ in how they actively remodel their microenvironment and sustain niche support. Rather than representing a single biological state, the persisting AML cells may therefore comprise multiple niche-associated LSC subtypes within the same leukemia, differing in quiescence, inflammatory signaling, and capacity for stromal or vascular remodeling. Therapeutic strategies targeting the LSC niche may need to account for this heterogeneity.

Together, our study supports a model in which LSC burden shapes the bone marrow niche through coordinated stromal and vascular remodeling. The dominant effect of LSC-high AML is a TGFβ-linked fibro-inflammatory shift of the stromal compartment, reinforced by CREB3L1 and potentially THBS1-dependent extracellular matrix feedback. In parallel, AML alters the vasculature through local infiltration-associated damage and condition-specific endothelial activation. These findings argue that AML niche remodeling is not only a consequence of overall disease burden, but is actively instructed by specific ligand-expressing LSC states. Therapeutically, this suggests that targeting LSC-supportive niche remodeling, particularly the TGFβ–CREB3L1 axis and its downstream extracellular matrix feedback, may weaken microenvironmental support for LSC persistence.

### Limitations of the study

The comparatively subtle transcriptional differences in the endothelial compartment recovered by single-cell profiling likely reflect, in part, a technical sampling bias: heavily infiltrated regions with structurally disrupted endothelium are less likely to yield viable single cells for droplet-based capture, biasing recovery toward less-infilitrated marrow. This may explain why endothelial changes were subtle by single-cell profiling despite clear vascular disruption by imaging.

Our findings rely on an immunodeficient xenograft model (NSGW41 mice). The KitW41 mutation permits robust AML engraftment without irradiation, avoiding a major confounding effect for a niche-based study, but the model lacks adaptive immunity and NK cells, so immune-niche crosstalk cannot be assessed.

A further limitation is that inferring cell-cell communication between human hematopoietic cells (HPSCs/AML) and mouse niche cell populations, which required converting human gene identifiers to their mouse orthologs prior to analysis (MultiNicheNet). Genes lacking a clear one-to-one mouse orthologue could not be included, and predicted interactions assume that orthologous ligand-receptor pairs retain equivalent binding efficiency and specificity across species, which might not always be the case.

## METHODS

### Human bone marrow and cord blood samples

Bone marrow specimens were collected from adult healthy controls and post-allo-HCT AML patients (**Supplementary Table 3**) after written informed consent, in accordance with the Declaration of Helsinki. Umbilical cord blood (CB) samples from healthy human infants were collected after written informed consent at the Department of Obstetrics at Heidelberg University. Sample collection and use for research were approved by the Research Ethics Boards of the Medical Faculty of Heidelberg University and EMBL Heidelberg (BIAC 2019-004 and BIAC Application No.: 2022-009).

CB samples were incubated for 10 min at room temperature (RT) with 10 µg/mL DNase I (Sigma-Aldrich, #DN25) and diluted 1:2 in CB-HSPC resuspension buffer (RB; 1% BSA, 10 mM EDTA in PBS). Mononuclear cells (MNCs) were isolated using a Ficoll-Paque Plus density gradient using SepMate tubes (15 ml Ficoll per tube; Thermo Fisher Scientific, #GE17-1440-02; STEMCELL Technologies, #85450), by centrifugation at 800 x g at RT (brake on). The MNC layer was collected, diluted in RB, and pelted (300 x g, 5 min, RT), and resuspended in RB. CD34^+^ hematopoietic stem and progenitor cells (HSPCs) were then isolated by immunomagnetic separation using CD34 MicroBeads (Miltenyi Biotec, #130-046-502). Per 3 x 10⁸ MNCs, 100 µl FcR Blocking Reagent and 100 µl CD34 magnetic microbeads were added, and cells were incubated for 30 min at 4°C. Cells were washed (diluted in 10 ml RB, 300 x g, 10 min, RT) and resuspended in 500 µl RB. The columns were primed twice with 500 µl RB, and CD34^+^ cells were magnetically retained and subsequently eluted in 1 ml RB after column removal from the magnet. Cells were frozen in freezing medium (10% DMSO in fetal bovine serum (FBS); Sigma-Aldrich, #F5724) at –80°C in a cooling container (−1°C/min).

### Xenotransplantation

NOD.Cg-*Kit^W-41J^Prkdc^scid^Il2rg^tm1Wjl^*/WaskJ (NSGW41) mice^16,17^ carrying a homozygous *Kit* mutation were used for xenotransplantation. Mice were bred and maintained in a specific pathogen-free facility under controlled environmental conditions (12 h light/12 h dark cycle, ambient temperature 18-21°C, relative humidity ∼55%) in individually ventilated cages at the German Cancer Research Center (DKFZ), Heidelberg. All animal experiments were approved by and performed in accordance with the guidelines of the Animal Care and Use Committees of the Regierungspräsidium Karlsruhe für Tierschutz und Arzneimittelüberwachung.

LSC-high (HLF wild type) and LSC-low (HLF KO) AML cells were generated from the patient-derived AML sample 04H112^13^ by CRISPR/Cas9 editing with single-stranded guide RNAs (sgRNAs) targeting GFP (sgGFP, control) and HLF (sgHLF), as previously described^12^. The patient-derived xenografted (PDX) AML cells were expanded by serial injections in NSGW41 mice. For each experiment, cells from three samples were pooled in saline (0.2 µm filtered), and 250K (imaging cohort) or 400K (scRNA-seq cohort) cells were injected intravenously in female (scRNA-seq cohort) or male (imaging cohort) mice aged 8 – 10 weeks (n = 4 per group). CB CD34^+^ HSPCs were injected with 40K (imaging cohort) or 70K (scRNA-seq cohort) cells per mouse (n = 4 per group). Engraftment was monitored by bone marrow aspiration and flow cytometry, and total bone marrow was collected at 8 weeks (scRNA-seq), 11 weeks (imaging cohort), or 12 weeks (3D imaging cohort).

For bone marrow isolation, bones were crushed in IMDM supplemented with 2% FBS and 100 µg/ml DNase I, filtered through a 30 µm cell strainer, and red blood cells were lysed using RBC lysis buffer (0.8% ammonium chloride, 0.08% sodium bicarbonate, 0.03% disodium EDTA). After washing with PBS, single-cell suspensions were obtained for downstream analyses. Genomic DNA was isolated from bone marrow and genome editing efficiency was assessed by Sanger sequencing of the amplified target region. HLF protein loss was confirmed by Western blot.

### Western blot

Cell pellets were thawed, centrifuged (1200 rpm, 5 min, RT), washed once with PBS, and lysed on ice for 30 min in RIPA buffer (Thermo Fisher Scientific, #89900) supplemented with protease inhibitor cocktail (Sigma-Aldrich, #11836170001). Lysates were cleared by centrifugation (13,000 rpm, 15 min, 4°C), and supernatants were transferred to pre-chilled tubes. Protein concentrations were determined using Bradford reagent (Bio-Rad) with BSA standards (Bio-Rad). Equal amounts of total protein were mixed with 4x NuPAGE LDS sample buffer (Invitrogen, #NP0007) and NuPAGE reducing agent (Invitrogen, #NP0004), and denatured at 95°C for 10 min.

Proteins were separated on 4-12% Bis-Tris gels (Invitrogen, #NP0322BOX; 80V for 10 min, followed by 120V for 1 hr) and transferred to nitrocellulose membranes (GE Healthcare, #10600003) at 100V for 1 hr at 4°C. Membranes were stained with Ponceau S to verify transfer, washed with PBST (0.1% Tween-20 in PBS), and blocked in 5% milk in PBST for 1 hr at RT. Primary antibodies against HLF (Abnova, #H00003131-M04, 1:500) and GAPDH (GeneTex, #GTX627408, 1:1000) were incubated overnight at 4°C in 5% milk in PBST. After three washes with PBST, membranes were incubated for 1 hr at RT with HRP-conjugated secondary antibodies (anti-mouse: Dianova, #115-036-062, 1:1000; **Supplementary Table 4**). Signal was detected using the SuperSignal™ West Femto substrate (Thermo Fisher Scientific, #34095) and imaged on an Amersham™ Imager 600 (GE Healthcare Life Sciences).

### Human bone marrow MSC isolation

Human bone marrow-derived mesenchymal stromal cells (MSCs) were isolated from bone marrow aspirates by plastic adherence *in vitro*, as previously described^76^. Briefly, mononuclear cells were isolated by Ficoll-Paque density gradient centrifugation and plated in MSC medium consisting of Dulbecco’s Modified Eagle Medium (DMEM; Thermo Fisher Scientific, #21885108) supplemented with 10% human platelet lysate (hPL heparin-free; PAN-Biotech, #P40-29050). Non-adherent cells were removed after 3 days, and adherent MSCs were maintained at 37°C and 5% CO₂ until ∼90% confluency (typically 1-2 weeks, passage 1). Cells were detached using 1x Trypsin (Sigma-Aldrich, #59427C) and reseeded at a density of 20,000 cells/cm². MSC identity and growth characteristics were consistent with previously established culture systems and phenotypic criteria. Expanded MSCs (passage 2) were subsequently used for *in vitro* co-culture experiments.

#### LSC-high vs LSC-low co-cultures for scRNA-seq

To profile MSC states under defined LSC burdens, the PDX-derived LSC-high and LSC-low AML cells were co-cultured with MSCs from four independent donors (post-allo-HCT AML patients). MSCs were seeded at 40,000 cells/cm² in 6-well plates and cultured to 80–90% confluency for two days. After two days, AML cells were seeded on top at a density of 20,000 cells/cm² and maintained in Stem Cell medium, containing IMDM (Thermo Fisher Scientific, #21980065) supplemented with 15% BIT (bovine serum albumin (BSA), insulin, transferrin; StemCell Technologies, #09500), 100 ng/ml SCF (Stem cell factor; Shenandoah, #100-04), 50 ng/ml FLT3-Ligand (Fms-related tyrosine kinase 3 ligand; Shenandoah, #100-21), 20 ng/ml IL-3 (Interleukin 3; Shenandoah, #100-80), 10 ng/ml G-CSF (Granulocyte-colony stimulating factor; Shenandoah, #100-72), 100 µM β-mercaptoethanol (Gibco, #21985023), 50 µg/ml Gentamicin (Thermo Fisher Scientific, #15750060), and 10 µg/ml Ciprofloxacin (GenHunter, #Q902-10ML). After 48 hrs, suspension and adherent fractions were collected separately and processed for FACS and scRNA-seq.

#### TGFβ perturbation of MSCs for bulk RNA-seq

To assess transcriptional responses to TGFβ pathway perturbation, MSCs from three independent old healthy donors were seeded in 12-well plates. Two days after seeding, the cells were treated with TGFβ1 (10 ng/ml; Thermo Fisher Scientific, #100-21-2UG), SB-431542 (10 μM; MedChemExpress, #HY-10431), AEBSF (250 μM; MedChemExpress, #HY-12821), or DMSO as vehicle control, prepared in MSC medium. After 3 days, the medium was replaced with fresh compound-free MSC medium. Cells were monitored and harvested at days 0, 1, 2, and 4 after media refreshment. Images were acquired, and cells were lysed in RLT lysis buffer supplemented with β-mercaptoethanol (Qiagen, #73404) and stored at –80°C. The RNeasy Micro Kit (Qiagen) was used for RNA isolation, following the manufacturer’s instructions. The bulk RNA-seq libraries were prepared by the Genomics Core Facility at EMBL Heidelberg following a SMARTseq-based protocol. Libraries were sequenced using Illumina NextSeq 2000.

#### Perturbation co-cultures for flow cytometry

To assess the functional consequences of stromal pre-conditioning on AML cell support, MSCs from one donor were seeded at 4-6K/cm^2^ density in 48-well plates. As described above, MSCs were treated for 3 days. Medium was replaced with compound-free stem cell medium, and 50K PDX AML cells were added for 4 days. Suspension and adherent fractions were collected separately (supernatant by pipetting and adherent layer by TrypLE (Gibco, #11538856) dissociation) and processed for flow cytometry.

### Single-cell RNA-sequencing

#### Mouse dissection and cell isolation for scRNA-seq

Bone marrow and bone-lining stromal cells were isolated from NSGW41 mice using a protocol adapted from Baccin et al.^7^, Gomariz et al.^8^ and Sood et al. (in preparation). Hips, femurs, and tibiae were dissected and cleaned of surrounding tissue. Bone marrow was flushed with RPMI 1640 (Sigma-Aldrich, #R8758) supplemented with 2% FBS (Gibco, #10270-106), and collected separately from the bone fraction. The remaining bone material was further processed to isolate the cells from the bone lining. The bones were mechanically crushed in RPMI/2% FBS and subsequently digested twice with 2 ml Digestion Buffer containing 2 mg/ml Collagenase IV (ThermoFisher Scientific, #17104-019) and 1 mg/ml Dispase II (Gibco, #17105-041) in HBSS (ThermoFisher, #14175-053) for 10 min at 37°C. Digestion was quenched with FACS buffer (PBS/2% FBS), and the digested fractions were centrifuged and pooled.

Both bone marrow and bone-lining cell suspensions were subjected to red blood cell (RBC) lysis using ACK lysing buffer (Gibco, #A1049201) for 10 min at RT, followed by quenching with FACS buffer. For the bone-lining fraction, mouse CD45^+^ cells were depleted using CD45 microbeads (Miltenyi Biotech, #130-052-301) and LS columns according to the manufacturer’s instructions. From the bone marrow fraction, cells were kept aside for final engraftment determination. For the rest of the bone marrow fraction, mouse CD45^+^ cells were depleted.

Next, cells were stained with antibody cocktails for FACS (**Supplementary Table 4**). Bone marrow suspensions were split and incubated with separate staining cocktails depending on the target populations: human HSPCs/AML cells or mouse HSPCs. Human and mouse hashing antibodies (TotalSeq-B, Biolegend) were included in the staining cocktails to enable sample multiplexing (**Supplementary Table 4**).

Before sorting, engraftment of human HSPCs/AML cells was confirmed for each mouse. Only mice with sufficient and equal engraftment were included; all mice in this cohort met this criterion. Mice within the same condition were pooled with equal cell numbers and processed together. FACS was performed with BD FACSAria Fusion Cell Sorter. Hematopoietic cell populations (10,000/condition) were sorted in 1.5 ml collection tubes containing 200 μl PBS/0.4% BSA, and niche populations (stromal (20,000/condition) and endothelium (4,000/condition)) were sorted into a V-bottom 96-well plate containing a droplet of 5 μl PBS/0.4% BSA per well. compartment. Humans and mouse hashing antibodies (TotalSeqB, Biolegend) were added directly to the FACS staining cocktail (**Supplementary Table 4**).

For single-cell RNA sequencing, the mouse HSPCs enriched from the bone marrow fraction and human hematopoietic fractions enriched from both the bone marrow and bone-lining fractions were pooled per condition. Cells were counted, and 16,000 cells were loaded in individual lanes per condition. The completely sorted niche fractions were loaded in individual lanes per condition. Single-cell RNA sequencing libraries were prepared using 10x Genomics Chromium Next GEM Single Cell 3′ Gene Expression v3.1 kit, according to the manufacturer’s instructions. Libraries were sequenced using Illumina NextSeq 2000.

#### Preparation of co-cultured human MSCs for scRNA-seq

Following 48 hrs of co-culture with LSC-high or LSC-low AML cells, the adherent stromal layer was detached using 1x Trypsin, and by FACS, the AML cells were depleted. The resulting huCD45^-^ alive fraction was collected for library preparation. Single-cell RNA-seq libraries were generated using the 10x Genomics Chromium Next-GEM Single Cell 3′ Gene Expression v3 kit. Libraries were sequenced using Illumina NextSeq 500.

### Immunofluorescence MSCs

MSCs cultured on poly-L-lysine (EMD Millipore, #A-005-C) coated coverslips were allowed to fix for 10 min at RT using 4% paraformaldehyde (methanol-free PFA; Thermo FIsher, #28908), then permeabilized using 0.1% TritonX (Sigma-Aldrich, #T8787-50ML) for 10 min at RT. Cells were blocked using 2% BSA for 45 min at RT and afterwards incubated with the primary antibody against CD90 (Abcam, #ab181469) for 1 hr at RT. After washing with PBS, cells were incubated with the secondary antibody (goat anti-mouse AF594, Invitrogen, #A-11032) for 45 min RT and mounted with Prolong Gold anti-fade mounting medium (Invitrogen, #P36966). Cells were imaged using an Olympus FV3000 confocal microscope.

### Histology

#### Tissue collection and embedding

For cryosections, femurs from the 4th mouse of each condition (scRNA-seq cohort) were collected, cleaned from connective tissue, and fixed in 4% PFA for 4-6 hrs. Bones were subsequently treated sequentially with 15% and 30% sucrose, embedded in O.C.T. medium (Tissue-Tek, #SA62550), and snap-frozen using isopentane (Sigma-Aldrich, #M32631), and stored at −80°C. Frozen blocks were cut into 5 μm sections using C35 blades (CellPath) and Kawamoto cryofilm^77^ using a cryostat (Leica CM1950).

For FFPE sections, femurs from a separate cohort of mice (imaging cohort) were collected and fixed in 4% PFA for up to 24 hours, followed by 10% EDTA-based decalcification. Bones were embedded in paraffin and cut into 10 μm sections using a microtome (Leica).

For 3D volumetric imaging, femurs from the imaging cohort were fixed in 4% PFA for 24 hours and decalcified in 10% EDTA for 2 weeks, as previously described (Kokkaliaris et al., 2020). Bones were embedded in 4% low-melting agarose (Sigma-Aldrich) and cut into 220 μm thick sections using a vibratome (Leica VT1200S).

#### H&E staining

FFPE sections were deparaffinized and rehydrated using the following solutions: xylene (2 x 10 min), 1:1 xylene:ethanol (3 min), 100% ethanol (2 x 3 min), 95% ethanol (2 x 3 min), 80% ethanol (3 min), 70% ethanol (3 min), and H₂O (2 x 20 s). Sections were then stained as follows: Mayer’s hematoxylin (3 min; Sigma, #MHS16), H₂O dips, bluing reagent (1 min; Epredia, #6769001), H₂O dips, alcoholic eosin (2 min; Sigma, #HT110116-500ML), and H₂O dips. Sections were mounted and imaged with a Zeiss Axioscan 7 using a Plan Apo 20x (NA 0.8) air objective with extended depth of field acquisition, using ZEN software.

#### Immunofluorescence on cryosections

Cryopreserved femurs sectioned onto Kawamoto’s type 3(16UF) film (SECTION-LAB, Co. Ltd., Japan) were stored at −80°C. Before staining, tape sections were cut to fit 8-well cell culture chambers (Ibidi). Sections were post-fixed with 4% PFA for 10 min at RT, then bleached with 4.5% H₂O₂ and 24 mM NaOH in PBS under white light for 45 min. Permeabilization and blocking were performed for 1 hr with 5% BSA in 0.1% PBS-Triton X-100. Primary antibodies (**Supplementary Table 4**) were diluted in 1% donkey serum (DS; Jackson ImmunoResearch) in 0.2% Triton X-100 and incubated overnight at 4°C. Secondary antibodies (**Supplementary Table 4**) were incubated for 1 hrs at RT in the dark. Nuclei were stained using DAPI (Thermo Fisher). Tape sections were mounted on microscopy slides with ProLong Gold Antifade Mountant (Invitrogen, #S36938), coverslipped, and sealed. Z-stacks were acquired with a Nikon Eclipse Ti2 Spinning disc confocal microscope equipped with 405 nm, 488 nm, 568 nm, 647 nm, and 750 nm solid state lasers, and an EMCCD camera (iXon3, Andor Technologies), using Plan Apo 20x air-(NA 0.8) and 40x oil-(NA 1.3) objectives.

#### 3D imaging of thick femoral sections

220 μm agarose-embedded femoral sections were blocked and permeabilized for 2 hrs (Tris-buffered saline, 20% DMSO, 0.05% Tween-20, and 10% DS (Jackson ImmunoResearch)) as previously described^78^. Primary antibodies against CD105, Laminin, human CD45, and Collagen type I together with Osteopontin were coupled with CF633, CF680, CF543, and CF750 secondary antibodies, respectively. Nuclei were stained using DAPI (Thermo Fisher). Sections were optically cleared in a step-wise manner (2,2’-thiodiethanol; Sigma-Aldrich) and imaged with a Leica Stellaris 8 equipped with two HyD-S, two HyD-X, and one HyD-R detectors and two laser lines (405 and white-light laser) using a 20x multiple-immersion objective (NA 0.75, FWD 0.680 mm) at 400 Hz, 8-bit and 1024 x 1024 resolution.

### Image analysis

H&E images were analyzed using QuPath software. Immunofluorescence images were analyzed using Fiji. Quantitative analysis of thick femoral sections was performed using Imaris (v9.9.0); for volumetric quantifications, regions with a size of 800 x 800 x 80 µm^3^ were selected and 32 sinusoid diameters were measured per region. For each tissue-wide scan, 2-4 regions were defined (control: 6 regions; 7%: 8 regions (4 low, 4 high); 47%: 6 regions (4 low, 2 high); 92%: 8 regions (4 moderate, 4 high)).

### Bulk RNA-seq analysis and differential gene expression analysis

Demultiplexed FASTQ files from perturbed primary human MSC samples were processed using an in-house Snakemake pipeline, including adapter trimming, read alignment, feature counting, and quality control. The resulting counts table was used to generate a SummarizedExperiment object (v1.36.0) for downstream analysis. Differential gene expression analysis was performed using DESeq2 (v1.46.0;^79^), retaining only genes with at least 10 counts across samples. P values were calculated using the Wald test.

### scRNA-seq preprocessing and quality control

Reads were aligned to the GRCh38 (v2020-A), mm10 (v2020-A-2.0.0), and barnyard (GRCh38 and mm10, v2020-A) reference genomes and quantified using *cellranger count* (10x Genomics, v3.0.1). After barnyard reference alignment, single cells were assigned to single species. The log-transformed number of mm10 reads relative to GRCh38 reads was calculated to detect doublets composed of cells of both human and mouse origin. Following doublets exclusion, we used the ratio of log-transformed GRCh38-mapped UMIs divided by the mm10-mapped UMIs to accurately assign single cells to either mouse or human origin. Cells with a ratio greater than zero were considered of human origin. Cells with fewer than 250 genes and more than 10% mitochondrial reads per cell were excluded from downstream analysis. After splitting the dataset based on the species-assigned barcodes, single-reference genome re-alignment was performed. The counts from the single-species alignments were used for further downstream analysis using Seurat v3^80^.

### Normalization, dimensionality reduction, and clustering

Following quality control and before dimensionality reduction, raw counts were normalized to account for the sequencing depth per cell and sample using *SCTransform*^81^. On the SCT assay (*SCTransform* output), Principal Component Analysis (PCA) was performed, followed by Uniform Manifold Approximation and Projection (UMAP) on the first 50 PCs. Ribosomal and mitochondrial genes were excluded from the variable features. Cells were then grouped into clusters using the Louvain algorithm, using FindClusters function. To define the clustering resolution of the aforementioned function, Clustree v0.4.4 was used (resolution = 1.5)^82^.

Marker genes of the unsupervised clusters were identified using Seurat’s *FindMarkers* function on the RNA assay. Genes detected in at least 50% of cells per cluster (min.pct=0.5) were considered. Differentially expressed genes between clusters were identified using the Wilcoxon Rank Sum test. Cell cycle score (Supplementary Figure 8B) was estimated with the Seurat’s *CellCycleScoring()* function.

To demultiplex the individual samples per condition based on the TotalSeq-B hashing antibodies, Seurat’s *HTOdemux* function with default parameters was used^80^.

### Trajectory analysis

To construct a pseudotime trajectory, we used *Monocle3*^34^ v1.3.7 and divided the fibroblastic– and osteo-lineages in two separate partitions. For each partition, we learned the principal graph using *learn_graph()* function with the following parameters (*close_loop* = FALSE; *minimal_branc_len* = 30; *nn.k* = 30). Afterwards, we ordered cells on the principal graph by randomly selecting 100 cells from the selected cluster of origin (*Cd34+* fibroblasts I for fibroblast trajectory and Osteo-adipo-CARs for osteo trajectory) and averaging pseudotime values after 1000 permutations.

### Gene regulatory network and GRaNPA analysis

The *pySCENIC* (v0.12.1) workflow^36,83^ was run using an in-house constructed Snakemake pipeline^84^. For gene regulatory network (GRN) inference, *GRNBoost2* algorithm from the Arboreto package was used^85^. SCENIC analysis was performed on the raw scRNA-seq data. For predicting the transcription factor (TF) regulons, mouse v9 motif collection was used, mm10 refseq-r80 10kb_up_and_down_tss.mc9nr.feather and mm10 refseq-r80 500bp_up_and_100bp_down_tss.mc9nr.feather databases from cisTarget (https://resources.aertslab.org/cistarget/). For downstream analysis, we constructed the final GRN using TFs identified in 90% of permutations (n=50) from the SCENIC Snakemake pipeline. GRaNPA v1.0.4 was used on the final GRN to predict the importance of TFs during transitions between cell types for fibro-trajectory using *GRaNPA_Analysis()* function (differentially expressed genes estimated with log2 fold-change cutoff of 2, *min.pct* = 0.1 and adjusted p-value < 0.1).

### Cell type decomposition using CIBERSORT

To estimate cell type localization in the spatial niches, we reused laser capture microdissection (LCM) data from Baccin et al.^7^, comprising bulk RNA-seq data from the various spatial bone niches. For the *runCIBERSORT*() function from the RNAMagnet v0.1.0 R package, we used *myAUC* to exclude genes between 0.3 and 0.7 from the LCM-seq data and used the remaining genes for downstream decomposition analysis. We excluded clusters with fewer than 10 cells and calculated the predicted fractions within the spatial niches.

### Gene set enrichment analysis (GSEA)

Functional analysis of DE genes was performed using *ClusterProfiler.* The ClusterProfiler’s functions applied (with default parameters) were *enrichGO* for Gene ontology enrichment analysis, *compareCluster* for KEGG pathway analysis, *enricherFunction* for Molecular Signatures Database MSigDB (Hallmark collection), and *enrichPathway* for pathway annotation from ReactomePA. The background gene set was defined as all the genes expressed in the dataset. P values were adjusted using Benjamini-Hochberg, and the cutoff was set to 0.05. For endothelial cell-specific functional enrichment analysis, gene set^86^ variation analysis (GSVA) on 615 EC-related gene sets selected from the MSigDB database^87^.

### Reference-based scRNA-seq projection

To project stromal populations from our mouse scRNA-seq to the human reference dataset^25^ and co-culture scRNA-seq on the stromal compartment from our mouse dataset, we used ProjecTILs^75^ v3.4.3 R package. We constructed a reference embedding based on the reference atlas (made with *make.reference()* function with *ndim=50* and r*ecalculate.umap=FALSE* parameters) using *Run.ProjecTILs()* function and plotted on the reference UMAP with *plot.projection()* function.

### Cell-cell communication analysis

To infer ligand-receptor interactions between cell types from scRNA-seq data, we used MultiNicheNet^43,44^ v2.1.0 R package. As senders, we defined the human HSPCs or LSCs, and either the mouse stromal or endothelial populations as receivers. As the underlying ligand-receptor database, we used *lr_network_mouse_allInfo_30112033.rds* ligand-receptor network and *ligand_target_matrix_nsga2r_final_mouse.rds* ligand target matrix. We ran MultiNicheNet with *“AML LSChigh vs AML LSClow”* and *“AML LSClow vs AML LSChigh”* contrasts. For the downstream analysis, we considered only cell types with more than 10 cells. We computed the fraction of expression in a sample-agnostic way using *get_frac_exprs_sampleAgnostic()* function with *min_sample_prop = 1* and *fraction_cutoff = 0.05*. For the gene set enrichment of the differentially expressed target genes, we used cutoffs of 0.25 log2 fold-change and adjusted p-value <0.05. To compute the final prioritization scores per contrast, we used *ligand_activity_down = FALSE* parameter. Afterwards, we filtered ligand-receptor pairs based on the curation effort parameter (*curation_effort > 3*) and recalculated prioritization scores, modifying prioritization weight of activity_scaled as follows: *prioritizing_weights[“activity_scaled”] * dplyr::coalesce(max_scaled_activity, 0)*. For the final selection of ligands, we took the top 50 ligands by prioritization score in each contrast and visualized them with their respective receptors.

## ACKNOWLEDGEMENTS

We are grateful to members from the Zaugg and Pabst groups for their valuable input, in particular Charles Dussiau. We thank the tissue donors for their contribution. We thank Irmela Jeremias for providing us with PDX-346 and PDX-602, and Dr. Kawamoto (Tsurumi University Japan) for providing us with the cryofilm and the sectioning starter kit. We thank Franziska Pilz and Andrea Kuck for their support with the mouse single-cell experiments. We thank the members of EMBL’s Flow Cytometry Core Facility (FCCF), Genomics Core Facility (GeneCore), and Advanced Light Microscopy Facility (ALMF), as well the Single-Cell Open Lab (scOpenLab) and Flow Cytometry Core Facility at DKFZ for technical support for the single-cell experiments, sequencing, and imaging. We thank EMBL IT for providing the infrastructure and support in performing the data analysis. Schematics throughout the manuscript were made with BioRender (<u>biorender.com</u>).

## FUNDING

This work was supported by the Deutsche Forschungsgemeinschaft (DFG, German Research Foundation; SFB 1709/1 2025, 533056198) to J.B.Z., C.P., M.G.A.E., K.K., and C.M.T.; the Max-Eder-Grant of the German Cancer Aid (70114435) to C.P.; the European Union (ERC, epiNicheAML, 101044873) to J.B.Z.; the SNSF (P2ZHP3_199669) and EMBO (538-2021) Postdoctoral Fellowships to K.D.P.; the DFG (TRR1278-C06, WA2837/8-1) and Carl Zeiss Foundation (Impuls, P2019-01-006), and the DFG under Germany’s Excellence Strategy (EXC 2051, 390713860) to C.W.; and EMBL Core Funding to J.B.Z. and S.K.S.

The Fritz Lipmann Institute (FLI) is a member of the Leibniz Association and is financially supported by the Federal Government of Germany and the State of Thuringia. This work is also in part supported by the Health + Life Science Alliance Heidelberg Mannheim and received state funds approved by the State Parliament of Baden-Württemberg (“MULTI-SPACE”). The data storage service SDS@hd is supported by the Ministry of Science, Research, and the Arts Baden-Württemberg (MWK) and DFG through grants INST 35/1314-1 FUGG and INST 35/1503-1 FUGG.

Views and opinions expressed are however, those of the authors only and do not necessarily reflect those of the European Union or the European Research Council. Neither the European Union nor the granting authority can be held responsible for them.

## AUTHOR CONTRIBUTIONS

K.D.P., A.M., C.P., and J.B.Z conceived the project and designed the study. K.D.P., A.M., and S.S. performed the single-cell mouse experiments. L.H. and S.G. performed CRISPR/Cas9 perturbations and mouse injections. A.M., I.B., and R.M. processed and analyzed the scRNA-seq data. K.D.P., J.R., and I.B. processed and analyzed RNA-seq data. K.D.P., A.M., Y.B., C.H., J.J-L., and D.H. performed *in vitro* (co-culture) experiments. K.D.P., A.M., C.H., and R.R. processed mice for flow cytometry and imaging. K.D.P. and A.M. performed 2D imaging. T.R. performed 3D tissue imaging. A.K. designed the Shiny app. C.W. provided key reagents. K.D.P., A.M., I.B., C.P., and J.B.Z. compiled and interpreted all the data. K.D.P. and J.B.Z. wrote the manuscript, incorporating text from A.M. and with input from I.B. and C.P.. C.M.T., S.K.S., K.K., M.A.G.E., C.P., and J.B.Z. supervised the project. All co-authors read and agreed on the manuscript.

## CONFLICT OF INTEREST

The authors declare no competing interests.

## DATA AND CODE AVAILABILITY

Raw count data for single-cell transcriptomic profiling (10x Genomics) and bulk transcriptomic gene expression data are available upon request through the corresponding authors.

Code used for the analyses of the bulk and single-cell gene expression data are available upon request.

---

**Supplementary Figure 1:**
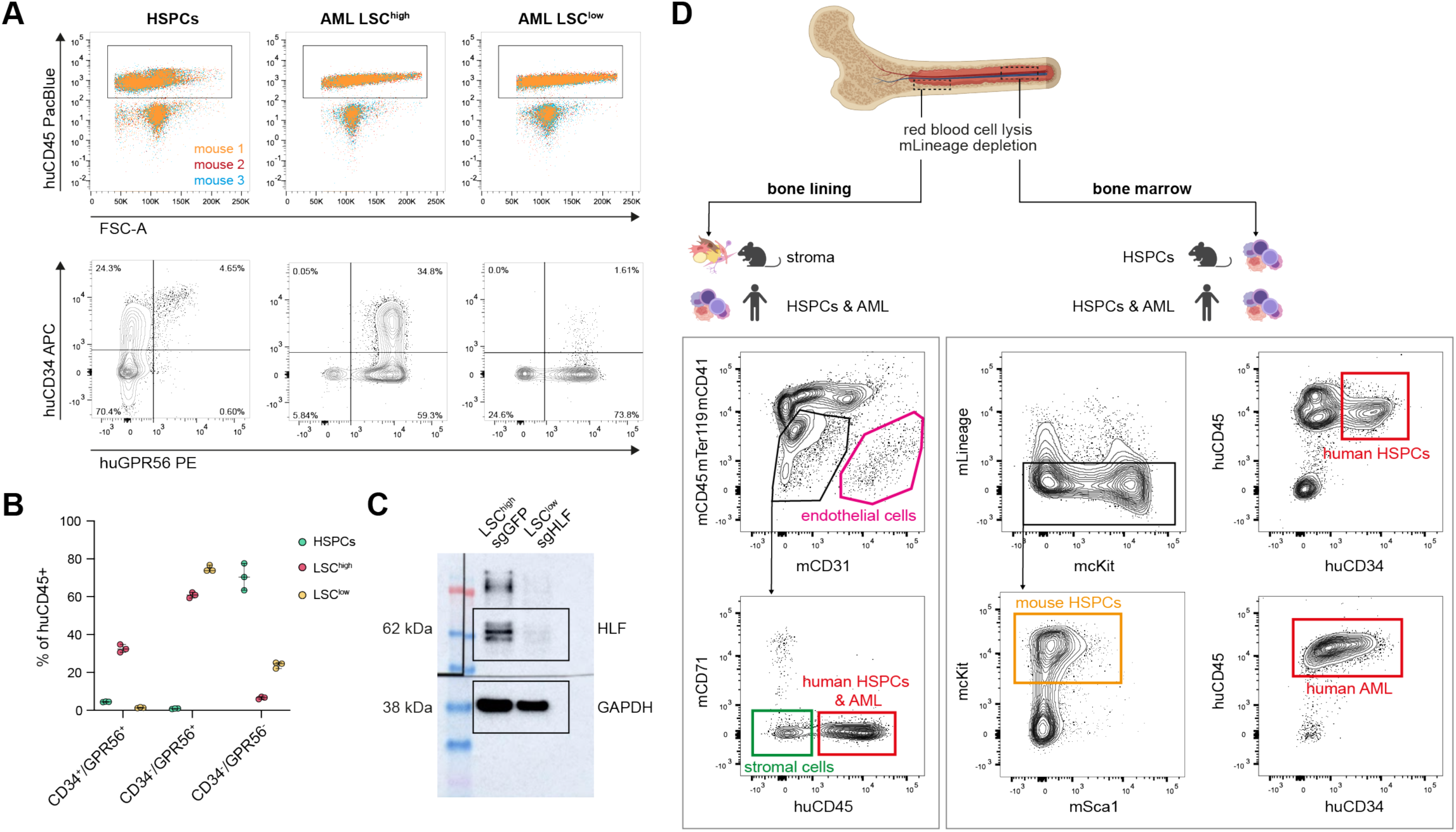
Flow cytometric definition and isolation of mouse niche, human HSPCs, and AML populations in NSGW41 xenografts. (A) Representative flow cytometry gating strategy for human hematopoietic stem and progenitor cells (HSPCs) and LSC-low and LSC-high AML populations in xenografted mice. Human cells were first gated based on huCD45 expression and forward scatter (FSC-A). Subsequent gating on huCD34 and huGPR56 distinguishes CD34+/GPR56+ (slow-cycling LSCs), CD34-/GPR56+ (fast-cycling LSCs), CD34-/GPR56-(blasts) fractions. Quadrant frequencies(% of huCD45+) are indicated. Individual mice (n = 3) are color-coded. (B) Quantification of CD34+/GPR56+,CD34-/GPR56+, and CD34-/GPR56-fractions(% of huCD45+) from HSPCs (green), LSC-high (pink), and LSC-low (yellow) AML. Data points represent individual mice (n = 3) across the conditions; bars indicate median with 95% confidence interval. (C) lmmunoblot of HLF protein expression in LSC-high (sgGFP = control sgRNA) and LSC-low (sgHLF = HLF KO) AML cells, confirming loss of HLF upon sgHLF; GAPDH serves as a loading control. Molecular weight markers (kDa) are indicated. (D) Schematic shows cell enrichment strategy of the mouse and engrafted cells for scRNA-seq. After red-blood-cell lysis and mouse lineage depletion, the flushed bone marrow fraction and digested bone lining fraction were stained for FACS. The bone marrow and bone lining fractions were processed separately. Endothelial cells (mCD31+), stromal cells (mlineage-, mCD71-, huCD45-), mouse HSPCs (mcKit+/m-Sca1+/-), human HSPCs (huCD45+/huCD34+), and human AML cells (huCD45+/huCD34+/-) were enriched. For sequencing, endothelial (5,000) and stromal (20,000) cells were pooled per condition into one 10x Chromium lane, and the hematopoietic fractions (mouse HSPCs, human HSPC/AML bone marrow, human HSPC/AML bone lining) were pooled into a separate 10x Chromium lane per condition.

**Supplementary Figure 2:**
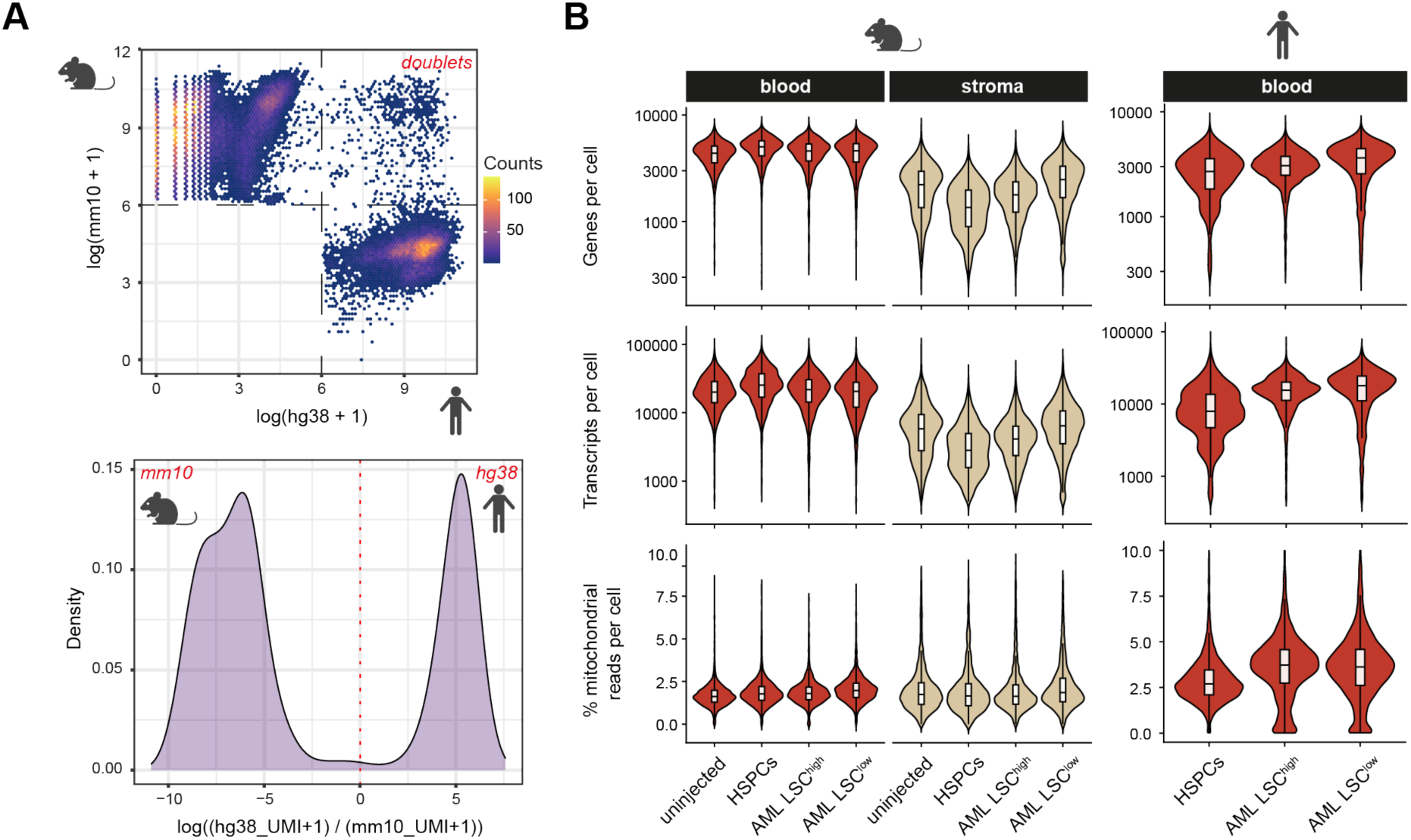
Species assignment, doublet exclusion, and quality control of scRNA-seq data from NSGW41 AML xenografts. (A) Species assignment and doublet identification based on transcript alignment to the human (hg38) and mouse (mm10) genomes. Top: density scatter plot of log(mm10 UMI + 1) versus log(hg38 UMI + 1) per cell barcode (color= cell counts). Cells with high expression from both genomes were classified as interspecies doublets and excluded from further analysis. Bottom: density distribution of log(hg38 UMls + 1) relative to log(mm10 UMls + 1). The bimodal distribution separates mouse from human cells, with the dashed line marking the threshold used for species assignment. (B) Per-cell quality metrics, with genes per cell (top), transcripts per cell (UMls; middle), and percentage of mitochondrial reads (bottom), shown as violin plots (overlaid box plot shows the median+ interquartile range) across the conditions, separated by the species (mouse or human cells) and enriched compartment (blood or stromal cell types). Metrics were used to define filtering thresholds for downstream analyses.

**Supplementary Figure 3:**
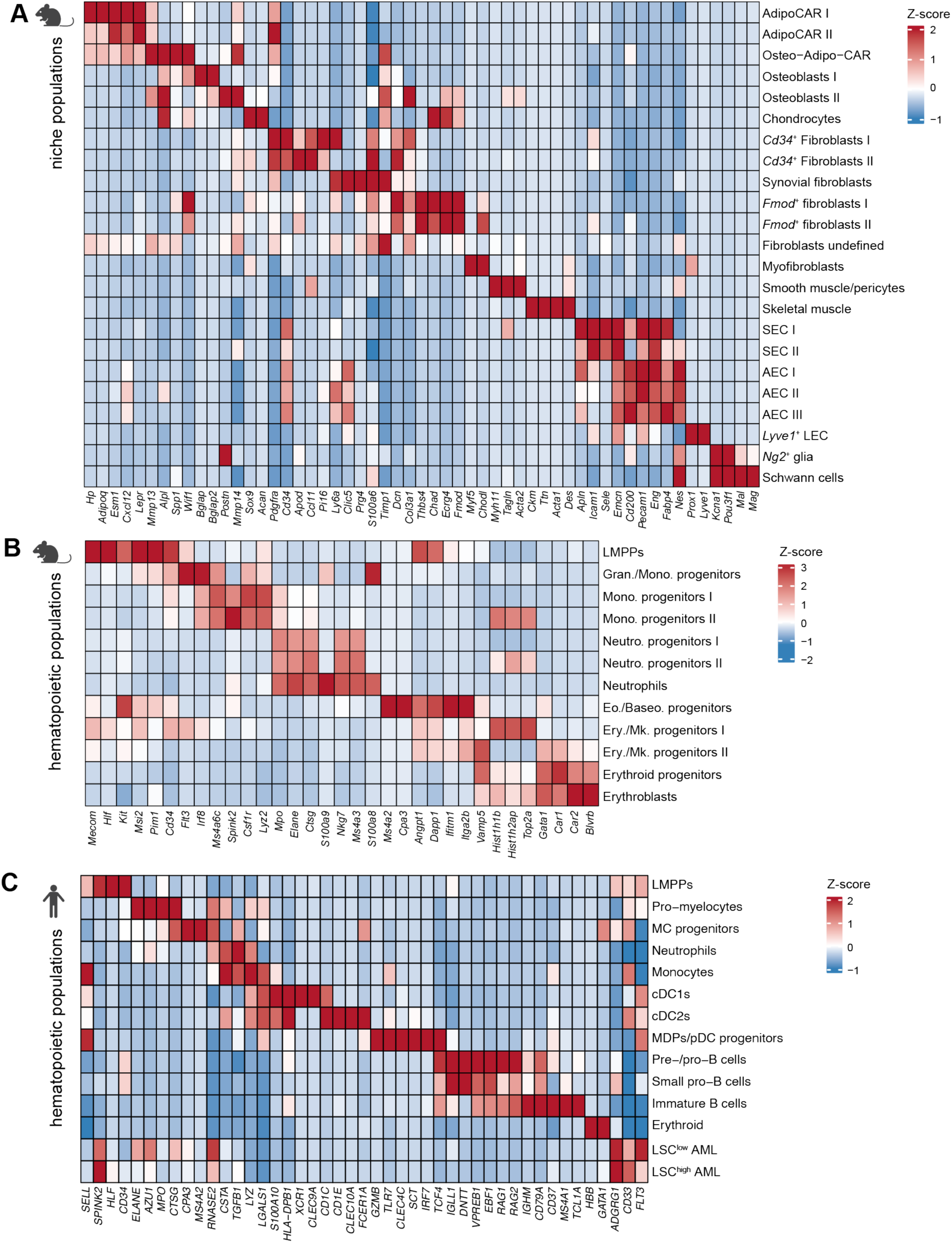
Transcriptional characterization of stromal and hematopoietic subpopulations in NSGW41 AML xenografts. (A) Heatmap showing scaled average expression of canonical marker genes across annotated mouse niche populations. Genes were selected based on literature-supported lineage markers and cluster-specific differential expression. (B) Heatmap showing scaled average expression of lineage-defining genes across annotated mouse HSPC populations. (C) Heatmap showing scaled average expression of marker genes across refined human hematopoietic clusters (CD34+ HSPCs-derived) and AML populations (LSC-low and LSC-high AML). For all panels, values represent gene-wise Z-scored average expression across annotated cell populations.

**Supplementary Figure 4:**
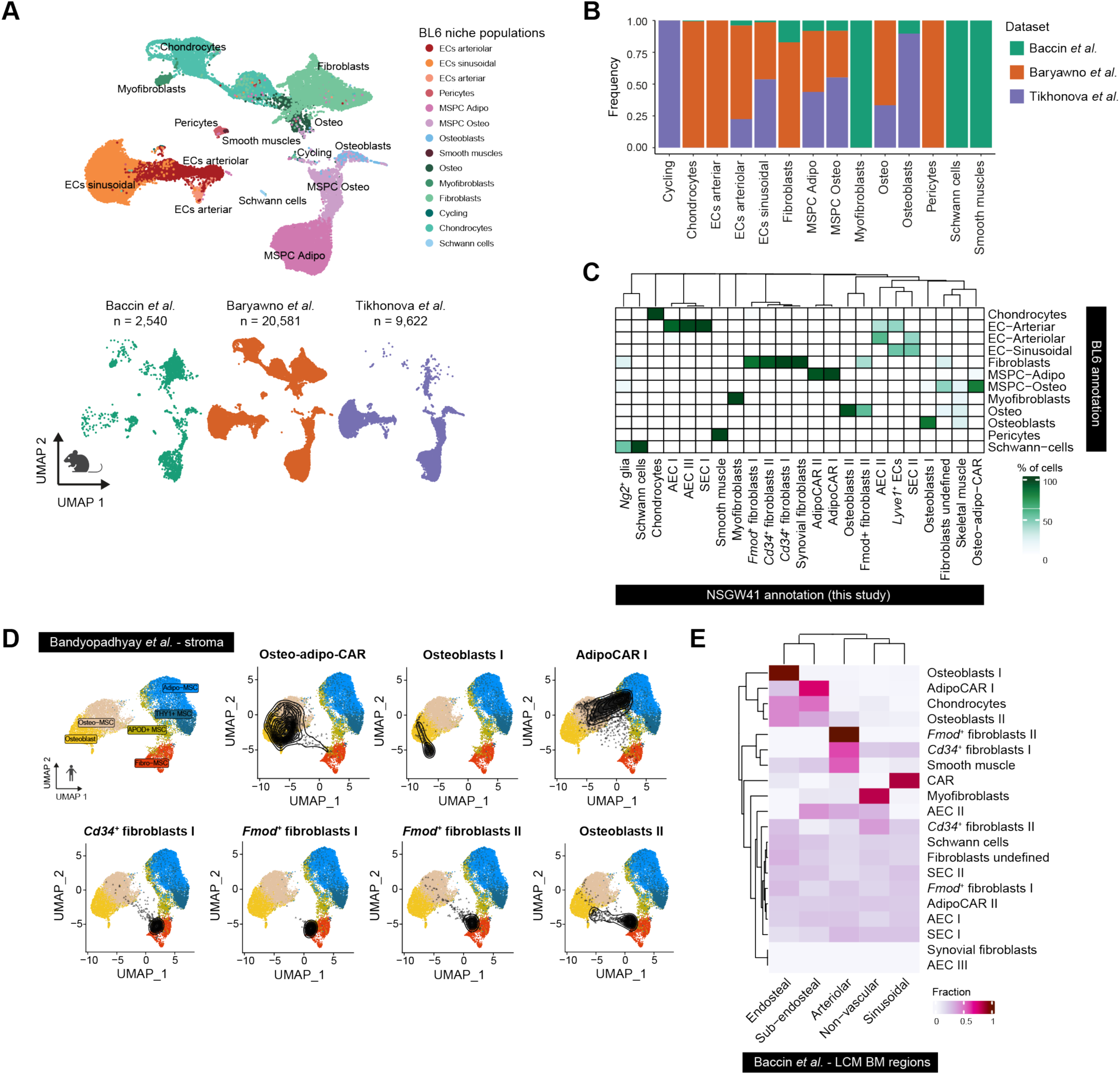
Cross-dataset comparison and spatial mapping of bone marrow niche populations. (A) UMAP visualization of integrated stromal and endothelial niche populations from C57BU6 (BL6) mouse bone marrow, annotated into major cell types (arterial and sinusoidal endothelial cells, pericytes, fibroblasts, mesenchymal stromal progenitors (MSPC Adipo and MSPC Osteo), osteoblasts, chondrocytes, myofibroblasts, cycling cells, Schwann cells)(Dolgalev and Tikhonova, 2021). Lower panels show the contribution of individual BL6 reference datasets (Baccin et al., Baryawno et al, Tikhonova et al.) to the integrated embedding. Baccin et al. applied a comparable digestion and cell enrichment strategy to our study(Baccin et al., 2020). Tikhonova et al. enriched BM nice cell populations using reporter lines(Tikhonova et al, 2019), and Baryawno et al enriched for stroma and endothelium from both flushed marrow and digested fractions(Baryawno et al., 2019). (B) Dataset composition of the annotated BL6 BM niche populations across the three reference datasets. Bar plots indicate the proportional contribution of cells from each dataset to individual cell type annotations, highlighting the differences in cell population representation across datasets. (C) Cross-annotation comparison between BL6 reference niche populations and NSGW41 niche populations from this study. The heatmap shows the percentage overlap of cells assigned to each population after label transfer and cluster correspondence analysis. Hierarchical clustering reflects transcriptional similarity across annotations. The color scale indicates the fraction of overlapping cells. (D) Projection of selected NSGW41 stromal populations onto the stromal reference embedding from healthy human bone marrow(Bandyopadhyay et al., 2024) using ProjecTILs(Andreatta et al., 2021). Contour overlays indicate the density of mapped NSGW41 cells within specific reference-defined regions, including CAR cells, osteoblast subsets, Cd34+ fibroblasts, AdipoCAR I population, and Fmod+ fibroblasts. (E) Spatial compartment assignment of NSGW41 stromal and endothelial populations. To annotate the single-cell niche population by BM location, we leveraged publicly available laser-capture microdissection (LSM) transcriptomic data from spatially defined regions(Baccin et al., 2020). Heatmap shows the resulting fractional assignments using CIBERSORT(Baccin et al., 2020) for each NSGW41 niche population (all conditions combined) across the five BM niches (endosteal, sub-endosteal, arteriolar, sinusoidal, and non-vascular niches). Hierarchical clustering groups cell populations with similar spatial compartment associations.

**Supplementary Figure 5:**
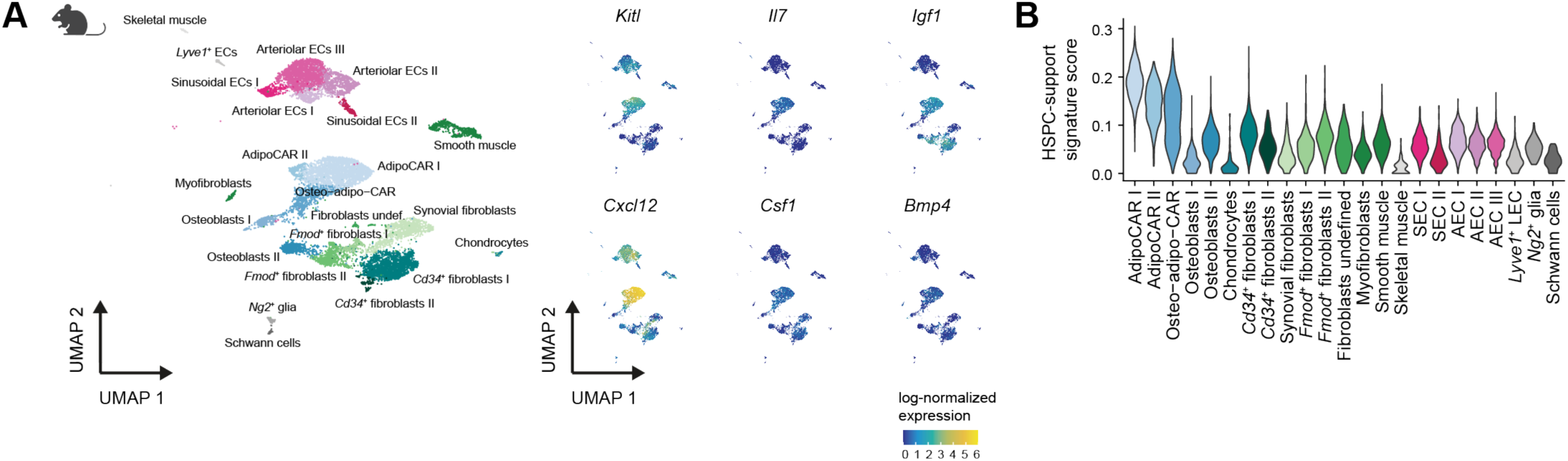
Hematopoietic-supportive programs in NSGW41 BM niche populations. (A) UMAP embedding of mouse BM niche populations (left) and feature plots (right) showing spatial expression patterns of selected key hematopoietic-supportive factors *(Kit/, 117, lgf1, Cxc/12, Csf1, Bmp4).* Color scale indicated log-normalized expression. (B) Violin plots showing HSPC-support signature score across stromal and endothelial populations. Scores represent per-cell enrichment and reveal preferential support capacity in specific stromal and endothelial subsets.

**Supplementary Figure 6:**
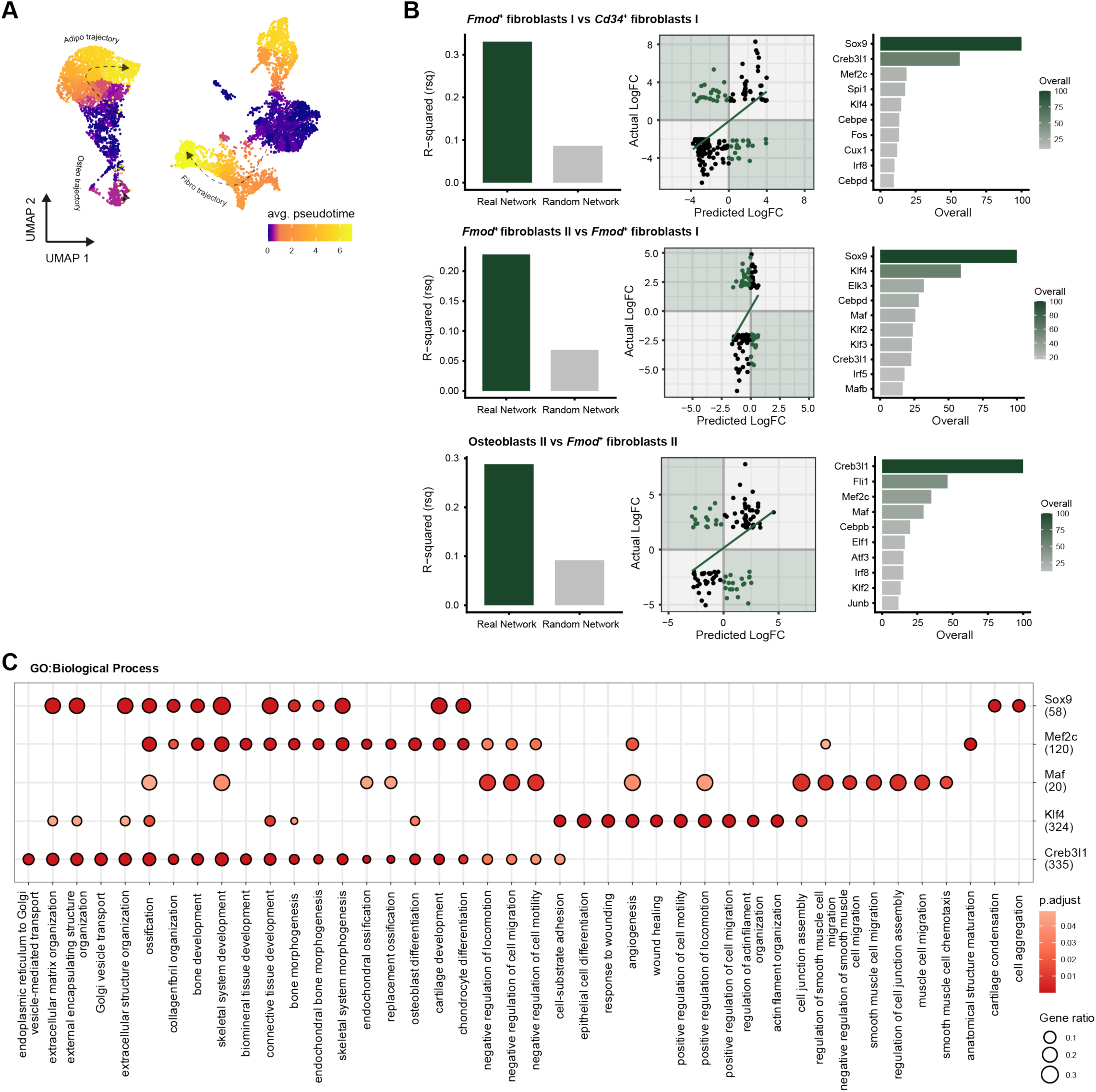
Stromal trajectory inference and gene regulatory analysis of state transitions. (A) Averaged pseudotime inferences across BM stromal populations using Monocle3. UMAP embedding shows inferred differentiation trajectories, including the CAR-derived adipogenic and osteogenic trajectories, and a fibrogenic trajectory extending from the stem cell-like *Cd34+* fibroblasts I through *Fmod+* fibroblasts states toward Osteoblasts II. Cells are colored by average pseudotime. (B) GRaNPA evaluation of TF-driven gene regulatory networks (GRN) for the three pairwise cell-state transitions. Left: model performance, shown as the variance in differential expression explained (R2) by the real GRN versus a randomized network control. Middle: GraNPA prediction accuracy, comparing the predicted with the actual log-fold-changes of differential expression (each dot= gene; line= linear fit). Right. top 10 most important TFs for each state transition, ranked by the GRaNPA importance score. (C) Gene Ontology (GO) Biological Process enrichment analysis of target genes associated with the top-five GRaNPA-ranked TF regulons along the fibrogenic trajectory, including Creb3I1, Klf4, Maf, Mef2c, and Sox9. Dot size indicates gene ratio relative to the tested gene set, and color indicates adjusted P value.

**Supplementary Figure 7:**
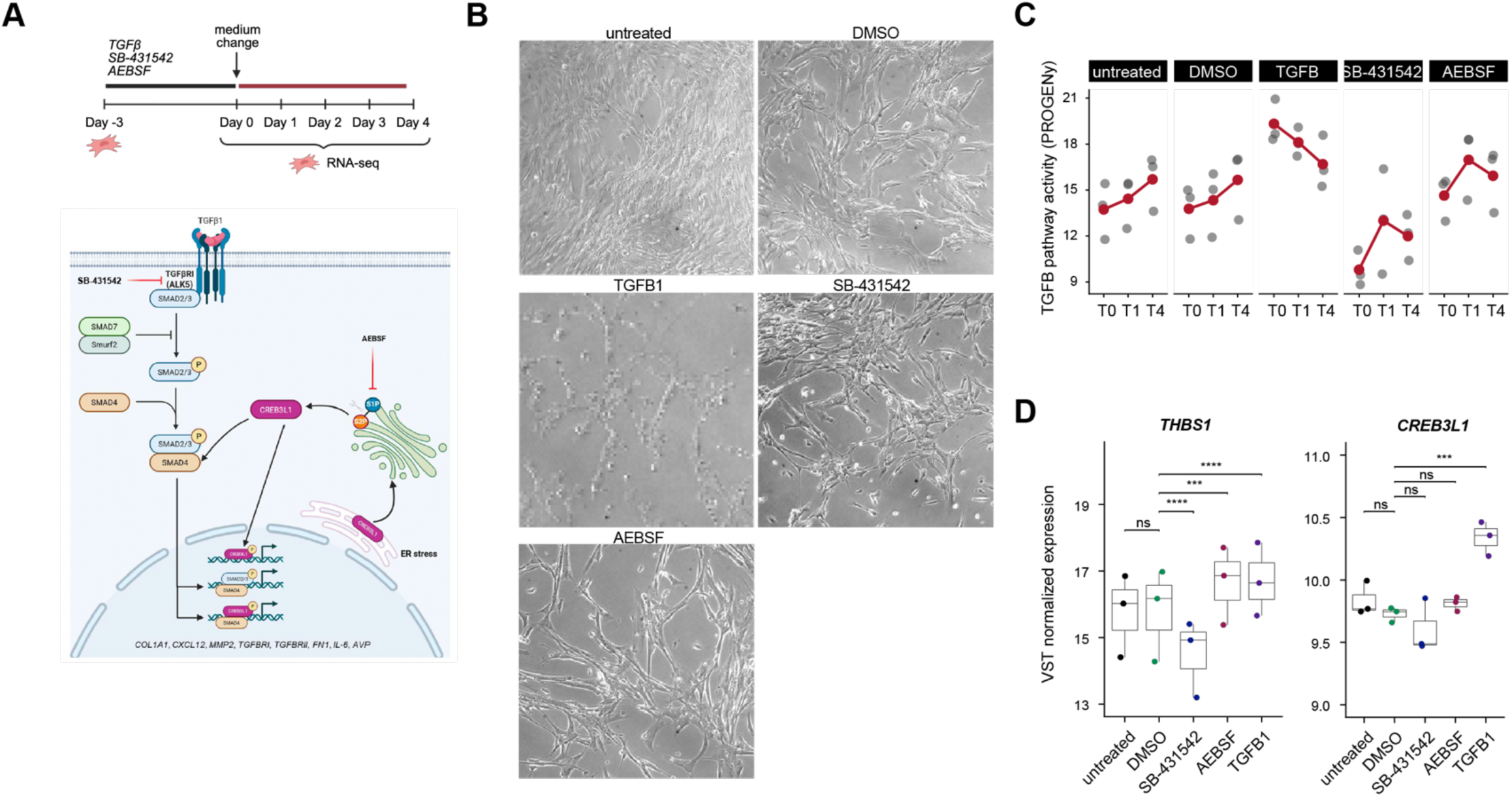
TGFl3 pathway perturbation in human bone marrow MSCs. (A) Top: experimental design for TGFl3 perturbation of human primary healthy BM MSCs. MSCs were pre-conditioned for 3 days (day –3 to day 0) with TGFJ31 (10 ng/ml), the TGFJ3 type I receptor kinase inhibitor S8-431542 (10 µM), the CREB3L1 deavage inhibitor AEBSF (250 µ M), or DMSO as vehicle control, followed by medium change and RNA collection at days 0, 1, and 4 for bulk RNA-seq. Bottom: schematic of the TGFj3--CREB3L1 signaling axis targeted by each perturbation. (B) Representative brightfield images of MSCs at day 0 following 3 days of treatment with the indicated conditions (untreated, DMSO, TGFJ3 1, SB-431542, AEBSF), illustrating condition-dependent morphological changes. (C) TGFJ3 pathway activity scores (PROGENy) across all five conditions at days 0 (TO), 1 (T1), and 4 (T4). Red dots indicate the median per time point; grey dots show individual donor values (n=3 donors). (D) VST-normalized expression of *THBS1* (left) and *CREB3L1* (right) across conditions at day 0. Each dot represents one donor (n=3); boxes show median and interquartile range. Adjusted P values from a DESeq2 Wald test are shown. –•p<0.005, •-•p<0.0001, ns = not significant.

**Supplementary Figure 8:**
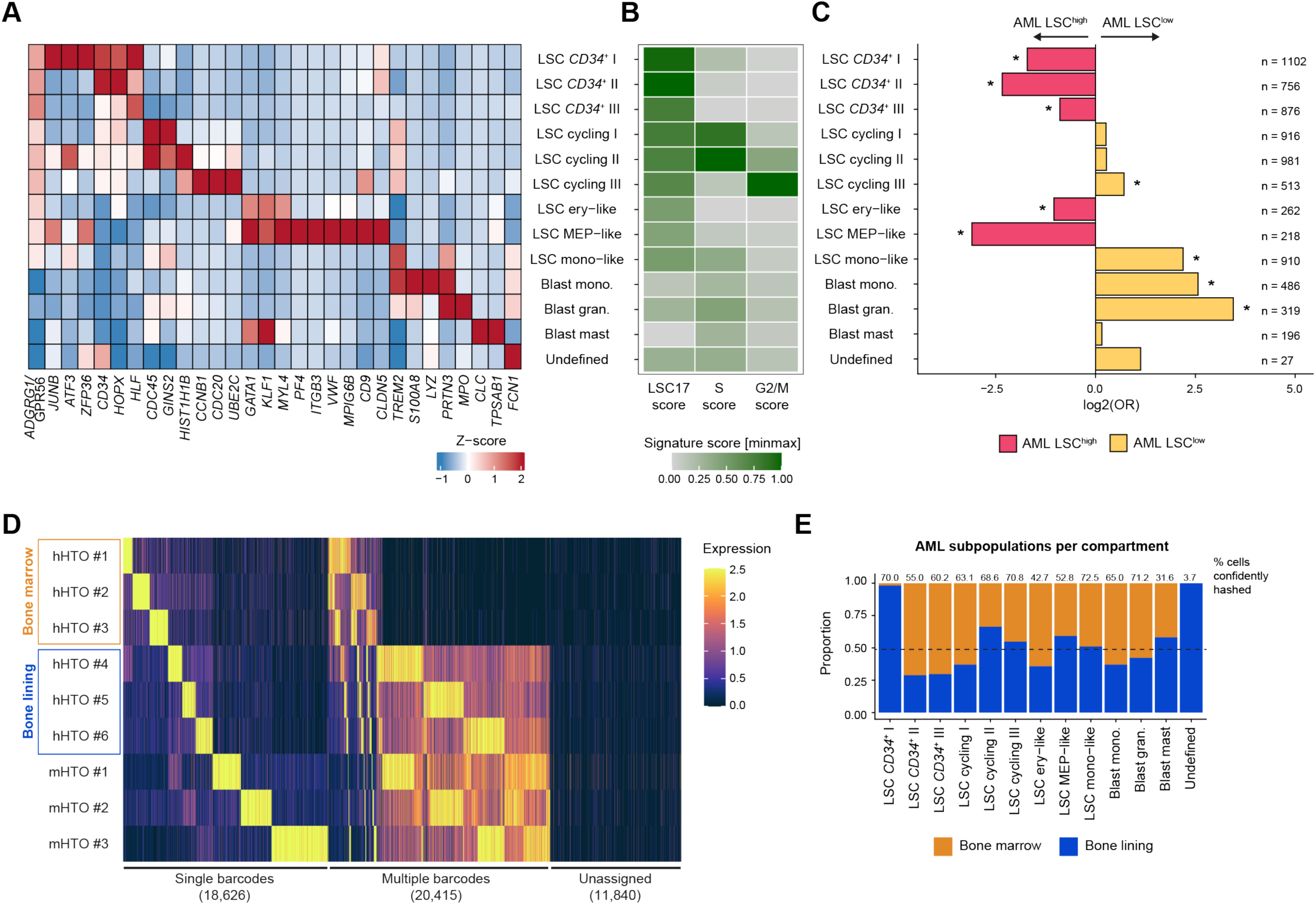
Subclustering and characterization of the human AML compartment. (A) Heatmap of marker gene expression (row-scaled z-score of mean expression) across the 13 annotated AML subpopulations, showing genes defining each cluster, including stem/progenitor, erythroid-megakaryocytic, cell cycle, myeloid, granulocytic, and mast markers. (B) Heatmap of signature scores (min-max scaled) for the LSC17 sternness signature (Ng et al., 2016) and the S-phase and G2/M cell-cycle scores. (C) Differential abundance of each AML subpopulation between LSC-high and LSC-low AML, expressed as the log2 odds ratio. The analysis was performed per cluster using Fisher’s exact test, adjusted p-value after Bonferroni correction are shown (*: <0.05). Values in pink indicate enrichment in AML LSC-high, while values in yellow reflect enrichment in AML LSC-low. n = number of cells per cluster. (D) Hashtag-oligo (HTO)-based demultiplexing used to assign cells to the bone marrow and the bone lining compartments (see also **Fig. 1A**, SFig. 1D, and **Methods).** Heatmap of HTO signal per cell for the human HTOs (bone marrow: hHTO #1-3; bone lining: hHTO #4-6) and mouse HTOs (mHTO #1-3). Cells were classified as carrying a single barcode, multiple barcodes, or unassigned. Cells confidently hashed (single barcode) were taken along for downstream analysis. (E) Compartment distribution (bone marrow: orange; bone lining: blue) of each AML subpopulation among hashed cells. Numbers above the bars indicate the percentage of confidently hashed cells per population. Dashed line markers 50%.

**Supplementary Figure 9:**
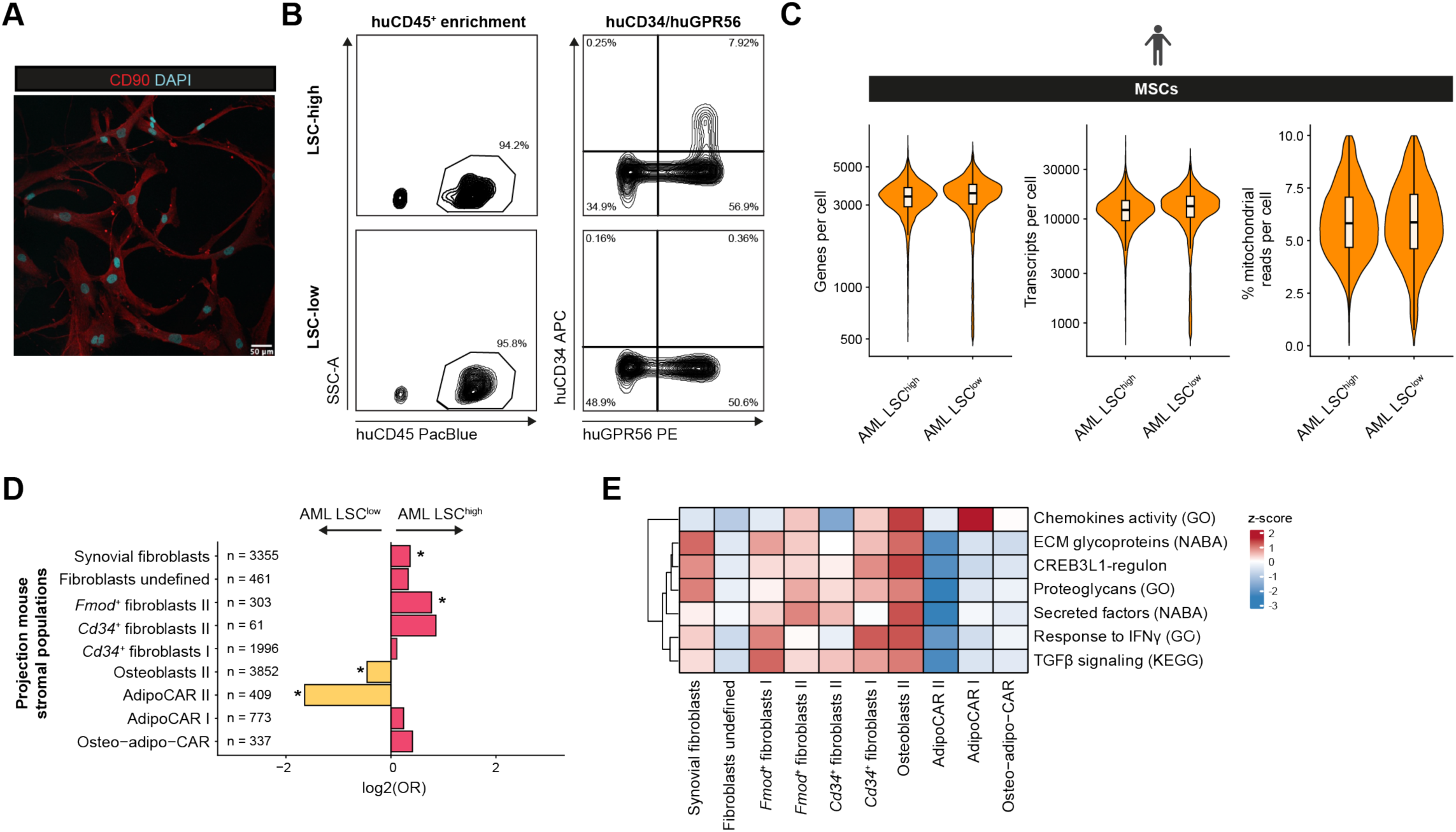
Human MSC co-culture with LSC-high and LSC-low AML. (A) Representative immunofluorescence image of human primary BM MSCs in co-culture, stained for MSC (CD90; red) and nuclei (DAPI; cyan). Scale bar: 50 µm. (B) Representative flow cytometry plots showing the gating strategy for AML enrichment from PDX models before co-culture. Left: human CD45+ AML cell enrichment by huCD45-PacBlue. Right subsequent gating on CD34 and GPR56 to distinguish the CD34+GPR56+ and CD34-GPR56+ AML fractions in LSC-high (top) and LSC-low (bottom) conditions. (C) Per-cell quality metrics for the scRNA-seq dataset shown as violin plots, comparing MSCs from LSC-high and LSC-low conditions: genes detected per cell (left), transcripts per cell (UMI count; middle), and percentage of mitochondrial reads (right). (D) Differential abundance of projected stromal subpopulations between LSC-high and LSC-low AML co-cultures, expressed as log2 odds ratio (Fisher’s exact test, Bonferroni-corrected, *: p<0.05). Pink bars indicate enrichment in LSC-high; yellow bars indicate enrichment in LSC-low. n = total cells per projected cluster across all donors. (E) Heatmap of average gene signature scores across projected human MSC subpopulations, shown as z-scores across rows. Signatures include chemokine activity (GO), ECM glycoproteins (NABA), CREB3L1 regulon, proteoglycans (GO), secreted factors (NABA), response to IFNy (GO), and TGFβ signaling (KEGG). Gene set sources GO, Gene Ontology; KEGG, Kyoto Encyclopedia of Genes and Genomes; NASA, Matrisome Project gene set categories(Naba et al. 2012).

**Supplementary Figure 10:**
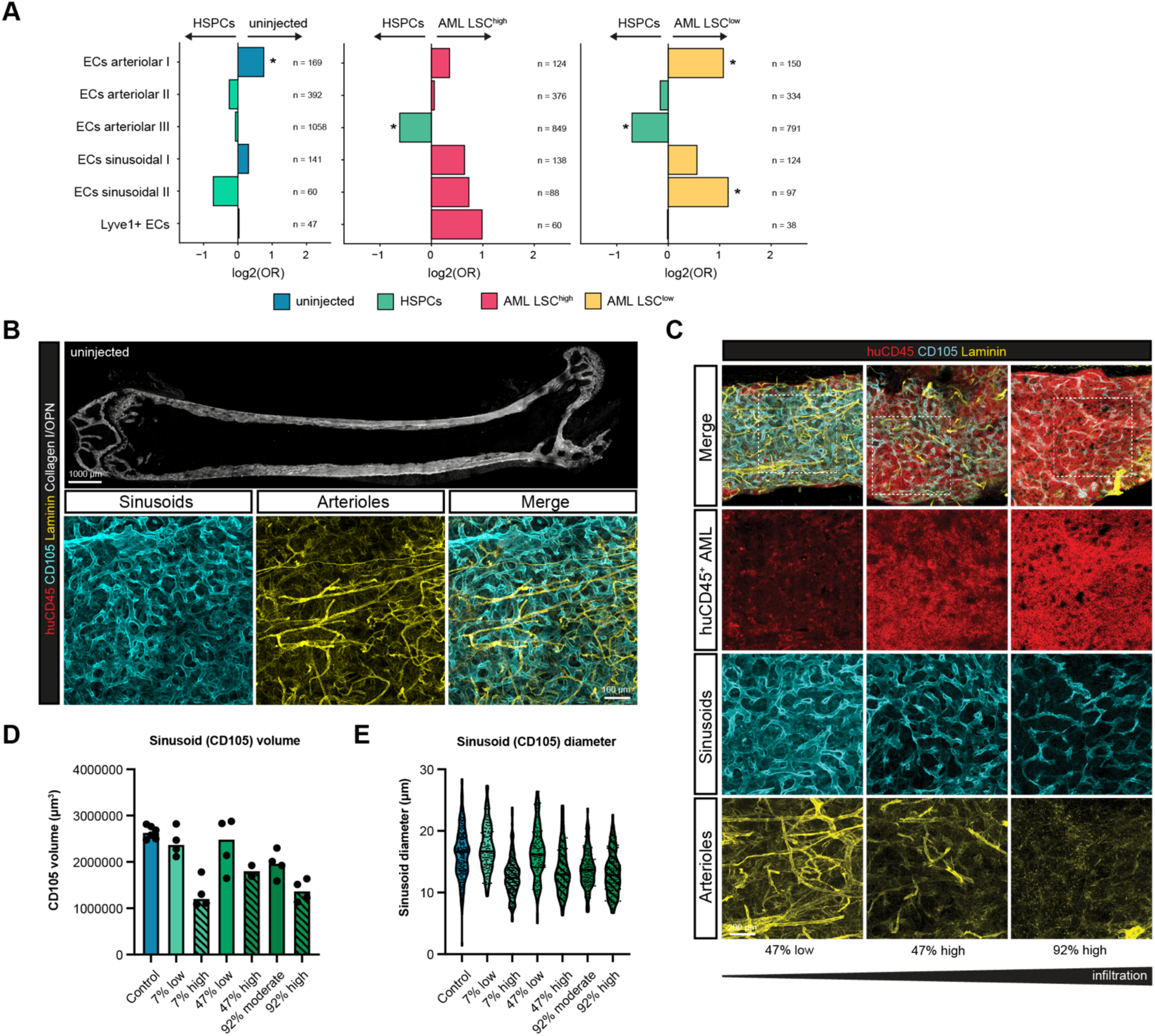
Endothelial compartment composition and bone marrow vascular architecture in NSGW41 xenografts. (A) Odds ratio (OR) plots showing relative abundance of endothelial subpopulations across pairwise comparisons: uninjected vs HSPCs (left), HSPCs vs AML LSC-high (middle), and HSPCs vs AML LSC-low (right). The analysis was performed per cluster using Fisher’s exact test, adjusted p-value after Bonferroni correction are shown (*: p<0.05). n = number of cells per cluster. (B) Representative image from volumetric (220 µm) tissue-wide immunofluorescence imaging of a femoral BM sample from an uninjected control mouse, stained for human CD45 (red; no human cells visible), CD105 (sinusoidal endothelium, cyan), Laminin (basement membrane/arterioles, yellow), and Collagen 1/OPN (bone surfaces, white). Upper panel shows the full femoral sample (scale bar: 1000 µm); lower panels show magnifications of sinusoidal (left) and arteriolar (middle) networks and their merge (right). Scale bar: 100 µm. (C) Representative magnified immunofluorescence images from AML-engrafted femoral sections stained for human CD45 (red), CD105 (cyan), and Laminin (yellow), shown for three regions of increasing leukemic infiltration: 47% low infiltration (left), 47% high infiltration (middle), and 92% high infiltration (right). Rows show the merged image (top), human CD45+ AML cells, sinusoidal endothelium (CD105), and arteriolar network (Laminin) separately. The gradient bar indicates increasing leukemic infiltration from left to right. Dashed boxes in the merge panel indicate the regions shown in magnified views below. Scale bars: 200 µm. (D) Bar plot showing CD105+ sinusoidal volume (µm’) per region across all individually quantified regions, grouped by mouse and infiltration level. Each dot represents one quantified region. Striped bars indicate high-infiltration regions; solid bars indicate low-infiltration or control regions. Full data underlying the paired analysis shown in **Fig. SH**. (E) Violin plot showing the distribution of individual sinusoid diameter measurements (µm) per region across all quantified regions (n = 30), grouped by mouse and infiltration level. Full data underlying the paired analysis shown in Fig. 51.

## Notes

### Competing Interest Statement

The authors have declared no competing interest.

